# Reconstituting human somitogenesis *in vitro*

**DOI:** 10.1101/2022.06.03.494621

**Authors:** Yoshihiro Yamanaka, Kumiko Yoshioka-Kobayashi, Sofiane Hamidi, Sirajam Munira, Kazunori Sunadome, Yi Zhang, Yuzuru Kurokawa, Ai Mieda, Jamie L. Thompson, Janet Kerwin, Steven Lisgo, Takuya Yamamoto, Naomi Moris, Alfonso Martinez-Arias, Taro Tsujimura, Cantas Alev

## Abstract

The segmented body plan of vertebrates is established during somitogenesis, a well-studied process in model organisms, but remains largely elusive in humans due to ethical and technical limitations. Despite recent advances with pluripotent stem cell (PSC)-based approaches^1–5^, a system that robustly recapitulates human somitogenesis in both space and time remains missing. Here, we introduce a PSC-derived mesoderm-based 3D model of human segmentation and somitogenesis, which we termed Axioloids, that captures accurately the oscillatory dynamics of the segmentation clock as well as the morphological and molecular characteristics of segmentation and sequential somite formation *in vitro*. Axioloids show proper rostrocaudal patterning of forming segments and robust anterior-posterior FGF/WNT signaling gradients and Retinoic Acid (RA) signaling components. We identify an unexpected critical role of RA signaling in the stabilization of forming segments, indicating distinct, but also synergistic effects of RA and extracellular matrix (ECM) on the formation and epithelialization of somites. Importantly, comparative analysis demonstrates striking similarities of Axioloids to the human embryo, further validated by the presence of the HOX code in Axioloids. Lastly, we demonstrate the utility of our Axioloid system to study the pathogenesis of human congenital spine diseases, by using patient-like iPSC cells with mutations in *HES7* and *MESP2*, which revealed disease-associated phenotypes including loss of epithelial somite formation and abnormal rostrocaudal patterning. These results suggest that Axioloids represent a promising novel platform to study axial development and disease in humans.

---

Previously, we and others were able to reconstruct the human segmentation clock *in vitro*^1,4,6^. However, these systems lacked the ability to form proper axial segmental organization –a central feature of all vertebrates– limiting their utility to understand how higher-order tissue organization and more advanced stages of human embryonic development occur. As supplementation of extracellular matrix (ECM) molecules *in vitro*, have been shown to facilitate the formation of higher-order tissue structures in organoids as well as help mimic morphogenetic processes in mouse pluripotent stem cell-derived gastruloids^2^ and trunk-like structures^3^, we set out to establish a single-germ layer, mesoderm-based 3D model of human axial development using human induced pluripotent stem cells (iPSCs) and ECM. iPSCs were exposed in a step-wise manner to signals promoting primitive streak (PS) and presomitic mesoderm (PSM) fates (Fig.1a and experimental procedures). The spontaneously symmetry-breaking and elongating mesodermal aggregates (Fig.1b, Extended Data Fig.1a, b) were then embedded into 10% Matrigel (MG), an ECM-rich culture supplement, which led to the spatiotemporally coordinated emergence of segments along the anterior-posterior axis of these growing and usually curved structures, which we termed Axioloids (Fig.1b, Extended Data Fig.1a-e). Axioloids were derived from two different human iPSC lines to ensure reproducibility and assessed for their morphological, molecular and functional features.

**Fig.1.**
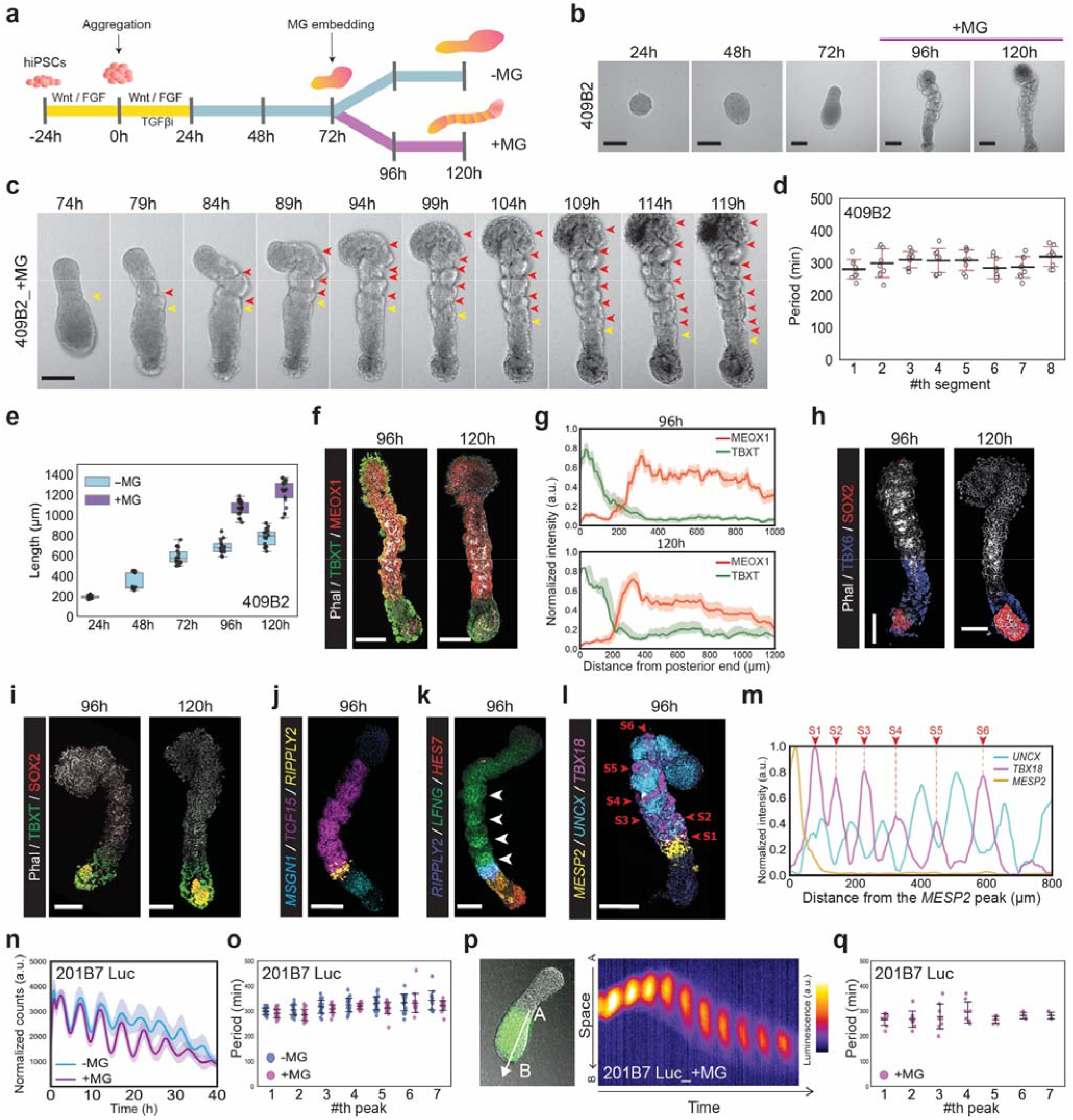
Generation of Axioloids from human pluripotent stem cells. **a,** Schematic summary of human iPSC-derived Axioloid induction protocol. Different color code for the major steps in the induction protocol: yellow, represents small molecule treatment, blue, suspension culture and purple, the Matrigel (MG) embedding phase; used abbreviations WNT, FGF and TGFβi stand for WNT agonist CHIR99421, basic FGF and TGFβ-inhibitor SB431542 respectively. **b**, and **c**, Time-lapse live imaging of human Axioloid induction. **b**, Representative bright field images of elongating Axioloids at 24h, 48h, and 72h, followed by images of Axioloids after MG embedding at 96h and 120h of culture. **c**, Serial images of a forming Axioloid at 5h intervals, from 74h to 119h (extracted from **Supplementary Video 1**). Colored arrowheads highlight the process of segment formation at each shown time point with the yellow arrowheads pinpointing areas where somite segmentation is ongoing, whereas red arrowheads highlight the areas where segmentation is completed. **d,** Periodicity of somite segmentation based live imaging observations (N=3, n=9). **e,** Axioloid length (antero-posterior) at 24h, 48h, 72h, and at 96h and 120h with or without MG embedding (N=4 with 24h, 48h and 72h n=18, and -MG 96h n=14 and +MG 96h n=24, and -MG 120h n=16, +MG 120h n=21). **f** and **g**, Immunofluorescence staining and signal quantification of MG embedded Axioloids at 96h and 120h. **f,** Merged channel images of Axioloids stained for F-actine (Phalloidin) in gray, TBXT (BRA) in green, and MEOX1 in red (corresponding single channel images are shown in Extended Data Fig.1i). **g,** Corresponding quantification along the posterior to anterior axis of TBXT (green line) and MEOX1 (in red) signal intensity (N=3, n=9). **h,** and **i,** Immunofluorescence staining of MG embedded Axioloids at 96h and 120h, images shown are representative of 3 independent experiments. **h,** F-actine (Phalloidin) is in gray, TBX6 in blue and SOX2 in red. **i,** F-actine (Phalloidin) is in gray, TBXT in green and SOX2 in red. **j** and **k,** HCR staining of MG embedded Axioloids at 96h, shown merged channel images are representative of 2 independent experiments. **j,** HCR staining of *MSGN1* in cyan, *TCF15* in magenta and *RIPPLY2* in yellow. **k,** HCR staining of *RIPPLY2* in blue, *LFNG* in green and *HES7* in red, white arrowhead highlight the strip pattern staining of *LFNG* in the posterior half of each somite. **l,** and **m,** HCR staining of MG embedded Axioloids at 96h for *MESP2* in yellow,*UNCX* in cyan and *TBX18* in magenta, and corresponding signal intensity measurement along the posterior to anterior axis normalized to the position of the *MESP2* signal peak. Numbered red arrowheads pinpoint the *TBX18* strip pattern observed in the posterior part of each somite. **n,** to **q,** Time series of a HES7: Luciferase expressing hiPSC cell line (201B7 Luc) from 72h to 112h. **n,** Quantification of the total HES7:luciferase signal in Axioloids with or without MG (N=2, n=4) and **o,** Periodicity was measured as the time interval between the consecutive HES7:Luciferase signal peaks (-MG N=6, n=22, +MG n=21). **p,** Right, kymograph along the line shown in the left image of Axioloid embedded in MG. **q,** Periodicity of the HES7:Luciferase signal measured in p, (N=4, n=8). Scale bar is 200μm.

## Axioloids display morphological and molecular features of the vertebrate embryonic axis and tail

To determine the similarities of Axioloids to the vertebrate embryonic axis and tail, we first assessed their morphological and molecular features. In Axioloids, segments appeared with a periodicity of about 4-5 hours (Fig.1c, d, Extended Data Fig.1f, g, Supplementary Video 1) and exposure to MG lead to a significant increase in convergent extension-based elongation, with Axioloid reaching total lengths of about 1000-1400 μm at 120h of culture (Fig.1e, Extended Data Fig.1h). These polarized axial structures could be further divided into a TBXT (also known as *Brachyury)* expressing posterior terminal domain, and an anterior MEOX1 positive somitic mesoderm (SM) region (Fig. 1f, g, Extended Data Fig.1i-k). Segments formed every 80-140 μm in the SM region of each Axioloid (Extended Data Fig.1l). The very end of the Axioloid tail was strongly positive for TBXT, with its expression reaching into and decreasing along the axis of the adjacent PSM, reminiscent of its reported expression in the tail bud (TB) and PSM of growing amniote embryos^7^. Interestingly, even in the absence of MG, this polarized pattern of expression of TBXT and MEOX1 could still be observed in Axioloids at 72, 96 and 120 hours of culture, suggesting that MG is not required for the initial establishment of this polarized expression pattern (Extended Data Fig.1m-q). Despite the regular nature of initial segment formation, fully epithelialized well-defined somite-like structures with proper apical-basal polarity and a central somitocoel-like cavity as seen in the developing embryo, were rarely observed in Axioloids (Extended Data Fig.2a, b). The segments in the MEOX1^+^ SM area of human Axioloids, which formed upon exposure to MG, were characterized by apical accumulation of actin, indicating the initiation of an epithelialization process (Fig.1f) but the epithelialization process within each segment appeared to be incomplete and largely disorganized (Extended Data Fig.2a, b).

Further assessment of gene and protein expression patterns of MG embedded Axioloids revealed a striking similarity to the anatomically and functionally regionalized molecular features described for the growing tail of vertebrate embryos. We could clearly distinguish a posterior-most TB region positive for TBXT and SOX2 (Fig.1h, Extended Data Fig.2c), neighboring with a presomitic mesoderm (PSM) domain clearly demarcated by regionalized expression of TBX6, *HES7* and *MSGN1* (Fig.1i-k, Extended Data Fig.2d-h). Further comparison of TBXT, SOX2 and TBX6 stained Axioloids revealed the presence of TBXT^+^ and SOX2^+^ double positive cells in the TB, which could be distinguished from TBX6^+^ but SOX2^-^ negative PSM cells (Fig.1h, i, Extended Data Fig.2c, d). We further found that the size of the TBX6^+^ PSM in Axioloids got reduced between 96h to 120h of Axioloid culture, while the SOX2^+^ population of the TB increased in size, which might be related to the observed slowing down of axial elongation and segment formation in Axioloids after 120h of culture (Fig.1h, i, Extended Data Fig.2c, d).

Besides the TB and PSM we could also distinguish a narrow *RIPPLY2* expressing anterior-PSM (aPSM) region (Fig.1j, k, Extended Data Fig.2e-g), which delineated a further rostrally localized MEOX1^+^ and *TCF15* expressing segmenting somitic mesoderm (SM) region, followed by a rostral-most MEOX1^+^ region of unsegmented mesoderm (Fig.1f, j, Extended Data Fig.2e, f). *LFNG (lunatic fringe)*, a major modulator of Notch signaling, was also expressed in the PSM, aPSM and the SM region of human Axioloids, with stripe-like expression pattern in SM similar to its expression reported for vertebrate embryos^8–10^ (Fig.1k, Extended Data Fig.2g, h). Emerging segments in the SM aspect of Axioloids, anterior to the *MESP2* expressing aPSM, showed also stripe-like expression of the rostrocaudal polarity genes *UNCX^11^* and *TBX18^12^*, indicating the establishment of proper anterior-posterior identity within forming segments of Axioloids (Fig.1l, m, Extended Data Fig.2i-l). Interestingly, *UNCX* and *TBX18* were expressed in the SM region of Axioloids even in the absence of MG, albeit disorganized and without a clear rostrocaudal pattern (Extended Data Fig.2m-p). Together, this demonstrates that Axioloids, albeit lacking a notochord and neural tube, share both morphological and molecular features of the vertebrate embryonic axis and tail.

## Axioloids recapitulate traveling-wave like oscillatory activity of the segmentation clock

A universal key feature of somitogenesis is the oscillatory activity of the segmentation clock, a molecular oscillator and gene regulatory network centred around Notch signalling active across the growing tail of the vertebrate embryo, and believed to control the pace and size of forming segments^13–15^. To determine dynamics of oscillatory activity within human Axioloids, we utilized a reporter system, which we had previously used to characterize the human segmentation clock *in vitro^1^*. We could clearly observe reproducible oscillatory activity of *HES7*, a well-studied segmentation clock gene^1,4,16,17^, with a periodicity of about 4-5 hours in Axioloids regardless of the presence of MG (Fig.1n, o). Furthermore, *HES7* displayed robust traveling-wave like expression from posterior-most TB and PSM region in an anterior direction along the PSM with a periodicity of 4-5 hours (Fig.1p, q, Extended Data Fig.2q-s, Supplementary Video 2). In contrast to earlier *in vitro* models of the human segmentation clock^1,4,6^, we observed the clear formation of segments in human Axioloids happening in striking synchrony with the segmentation clock. Segments formed every 4-5 hours in the anterior PSM region of Axioloids, a region, which overlaps and coincides with the wave front of *HES7* oscillatory activity (Extended Data Fig.2q-s, Supplementary Video 2). Our data thus indicates that both processes, segmentation and traveling wave-like oscillatory activity of the segmentation clock, are coupled in human Axioloids and happen in a spatiotemporally coordinated manner, a tight association that has so far only been reported in the embryonic tail of vertebrate model organisms^18,19^.

## Single cell RNA-seq analysis of human Axioloids

To further understand the functional features of our model system and to characterize the prospective dynamic changes in the cellular composition and molecular complexity of human Axioloids, we performed temporally-resolved single cell RNA-seq (scRNA-seq) analysis of various stages (48h, 72h, 96h & 120h) and conditions (+/-MG) of our system. Our analysis revealed the presence of multiple dynamically changing cell clusters, which could be matched to different mesodermal cell populations present in the developing axis and tail of vertebrate embryos, including tail bud (TB), presomitic mesoderm (PSM), anterior presomitic mesoderm (aPSM), somitic mesoderm (SM) and angioblast/endothelial-like (EC-like) cells (Fig.2a, b). These cell clusters could be further divided into putative subpopulations, based on their cell cycle state or time of emergence or respective maturation state during *in vitro* culture (Extended Data Fig.3a, b). RNA velocity analysis revealed a major differentiation trajectory, with early TB and early PSM cells dominating at 48h of culture, and giving rise to SM populations, which make up the majority of cells at 96h and 120h of culture (Fig.2b-d). Interestingly presence or absence of MG did not have a large effect on the differentiation trajectory or distribution of existing or emerging cell populations within Axioloids (Fig.2d, Extended Data Fig.3c). Many genes previously reported to be specifically expressed in the anatomical compartments and cell populations making up the mesodermal aspect of the embryonic tail, were identified and shown to also match the specific expression profiles in human Axioloids (Fig.2e, f, Supplementary Table 1). SM cells could be divided into six distinct clusters based on their expression profiles, with several of them showing specific high-level expression of various ribosomal proteins, including that of *RPL38*,for which mutations in mice were shown to result in homeotic transformations of the axial skeleton^20^ (Fig.2f, Extended Data Fig.3d). We observed that several genes, including *TBXT, CYP26A1, WNT3a* and *FGF8* that are initially highly expressed in early TB, showed reduced expression at later stages in culture, while *SOX2* remained strongly expressed in late TB. This suggests a possible change in number and function of putative somitogenic *TBXT* and *SOX2* double positive neuromesodermal progenitor cells (NMPs) present in the TB of human Axioloids, consistent with our earlier observation of decreased TBXT staining in SOX2^+^ TBs of 120h human Axioloids (Fig.2f, Extended Data Fig.3e, f, see also Fig.1h, i, Extended Data Fig.2c, d). RNA velocity and pseudotime analysis of scRNA-seq data from MG embedded, sequentially segmenting Axioloids at 96h of culture, revealed the presence of a major differentiation trajectory originating from TB and going over PSM and aPSM to SM cells, that matches with the posterior-to-anterior spatial distribution of these cell populations within human Axioloids (Fig.2g-i).

**Fig.2.**
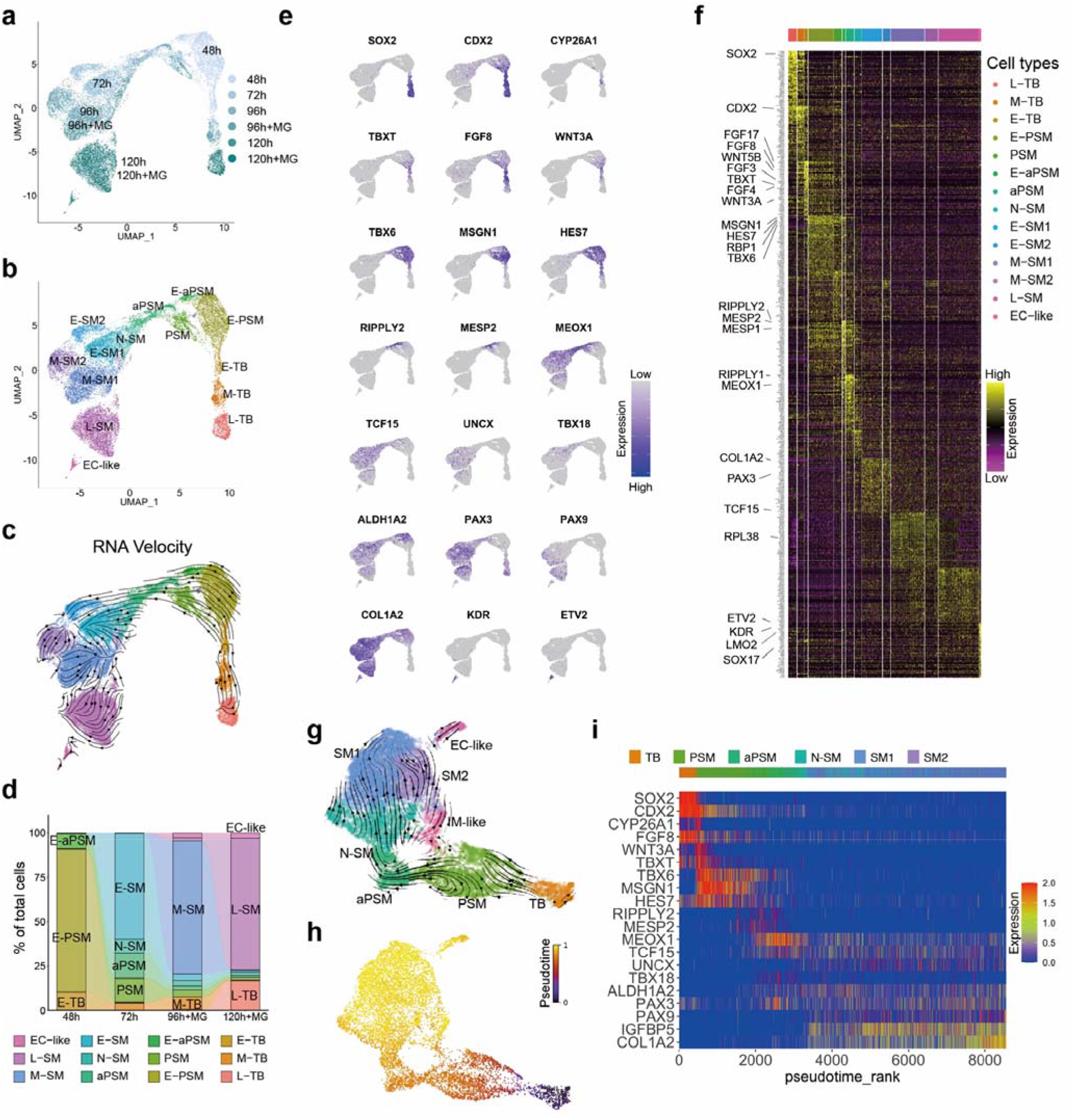
scRNA-seq characterization of human Axioloids. **a**-**c,** UMAP projection of scRNA-seq datasets of human Axioloids at 48h, 72h, 96h and 120h, colored by **a,** samples, and **b, c,** identified clusters. Both non-MG and MG samples are included for 96h and 120h. Arrows in **c,** show RNA velocity. **d,** Proportions of cell types in Axioloids over time. Regarding 96h and 120h, only MG-plus samples are shown. **e,** Expression levels of the selected genes are indicated on the UMAP plot. **f,** Single-cell expression profiles of identified marker genes for each cell cluster except for E-SM2, M-SM2, and presumably apoptotic cells. The top 50 (or less) genes of higher fold changes are shown. Presumably apoptotic cells are omitted. **g-h,** UMAP plots of Axioloids at 96h with MG, colored by identified clusters in **g** and by pseudotime in **h**. Arrows in **g** show RNA velocity. In this analysis, two replicates are integrated. **i,** Marker gene expression patterns at 96h with MG samples along pseudotime rank in **h**. IM-like and EC-like cells are omitted. Used abbreviations: TB (tail bud), E-TB (early tail bud), M-TB (mid tail bud), L-TB (late tail bud), PSM (presomitic mesoderm), E-PSM (early presomitic mesoderm), APSM (anterior presomitic mesoderm), E-APSM (early anterior presomitic mesoderm), N-SM (nascent somatic mesoderm), SM1 (somitic mesoderm 1), E-SM1 (early somitic mesoderm 1), SM2 (somitic mesoderm 2), E-SM2 (early somitic mesoderm 2), M-SM1 (mid somitic mesoderm 1), M-SM2 (mid somitic mesoderm 2), L-SM (late somitic mesoderm), IM-like (intermediate mesoderm-like), EC-like (endothelial cell-like), MG (Matrigel).

## Axioloids establish proper FGF/WNT gradients and express Retinoic Acid signaling components

As the embryonic tail of vertebrates is characterized by opposing gradients of FGF/WNT and Retinoic Acid (RA), believed to be involved in the establishment of the “wave front” during somitogenesis^21–23^, we next asked whether similar gradients are also present within our human Axioloids. Pseudotime analysis of our scRNA-seq data, which matched well with the anterior-to-posterior organization and spatial distribution of the major cell populations found in Axioloids, was used to predict the expression patterns of multiple FGF, WNT and RA signaling associated transcripts within Axioloids (Extended Data Fig.4a-c). HCR based *in situ* hybridization was used to validate the predicted spatial expression patterns of *FGF8* and *WNT3a*, two genes reported to be associated with the wave front during somitogenesis^24–26^, and revealed a clear posterior-to-anterior gradient of their expression in human Axioloids. Both genes were expressed strongest in the posterior TB and PSM region and their expression gradually decreased throughout the PSM up to the aPSM area marked by the expression of *MESP2* (Fig.3a, b, Extended Data Fig.4d-f). We then validated the expression of RA signaling associated molecules in Axioloids, focusing on *ALDH1A2*, an enzyme involved in the synthesis of RA from Retinal (RAL), and *CYP26A1*, a RA-metabolizing enzyme. We detected high level localized expression of *CYP26A1* in the TB, while *ALDH1A2* expression was high in the SM region of human Axioloids, showing highest expression rostral to the aPSM specific expression of *RIPPLY2* and slowly decreasing along the posterior to anterior axis (Fig.3c, d, Extended Data Fig.4g-i). Furthermore, *ALDH1A2* also formed a stripe-like expression pattern, showing higher levels of expression in the rostral halves of forming segments, similar to that described in the mouse^27^ (Fig.3c, d, Extended Data Fig.4g-i).

**Fig.3.**
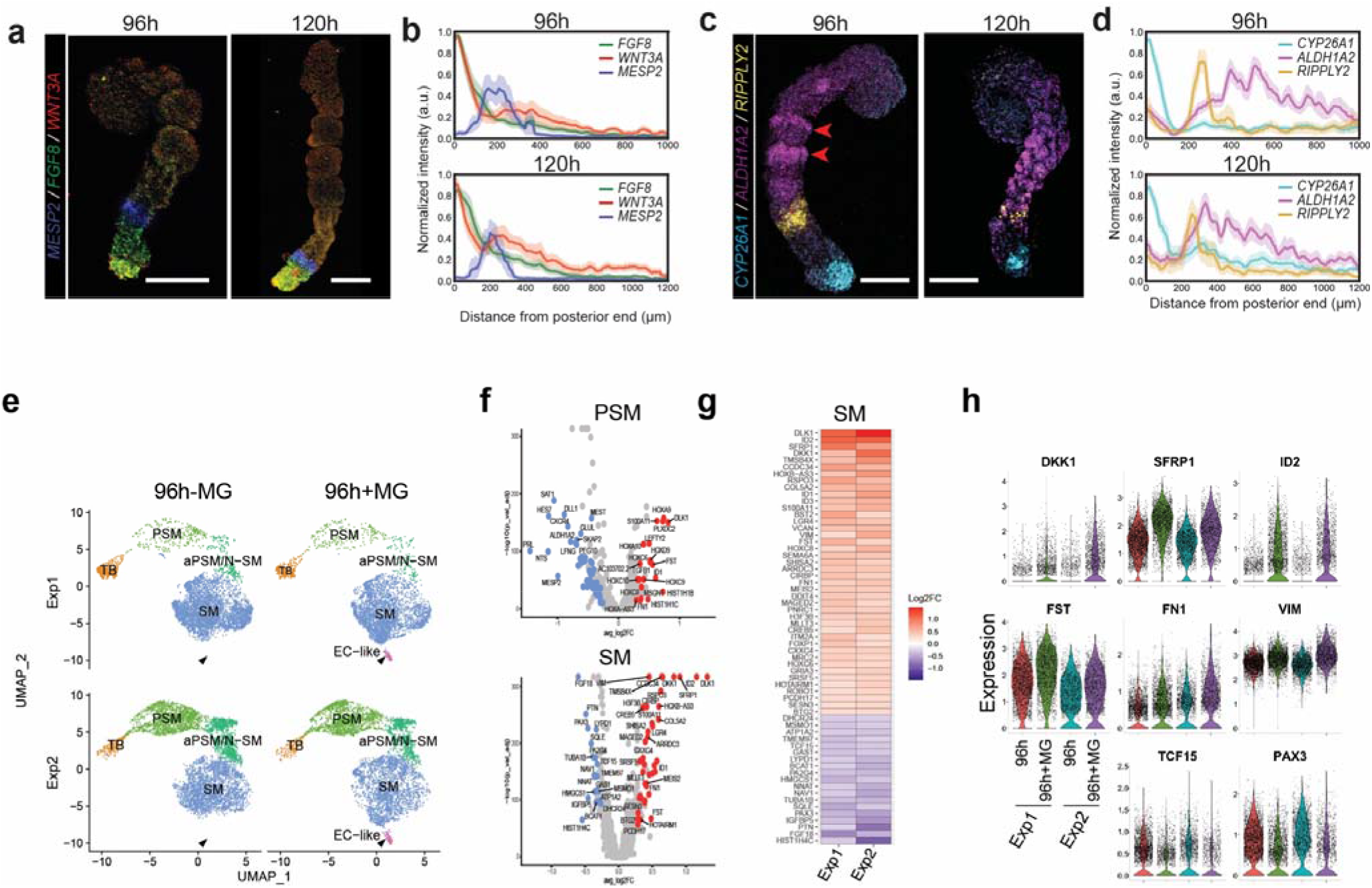
Signaling pathways and MG effect in Axioloids. **a-d,** Merged HCR staining images and signal quantification of MG embedded Axioloids at 96h and 120h. Shown images are representative of at least 3 independent experiments. **a,** HCR staining of *MESP2* in blue, *FGF8* in green and *WNT3A* in red **b,** corresponding quantification along the posterior to anterior axis of the signal intensity normalized the position of the *WNT3a* signal peak (96h: N=3, n=9 and 120h: N=4, n=10). **c,** HCR staining of *CYP26A1* in cyan, *ALDH1A2* in magenta and *RIPPLY2* in yellow and **d,** corresponding quantification along the posterior to anterior axis of the signal intensity normalized the position of the *CYP26A1* signal peak (96h: N=3, n=8 and 120h: N=4, n=9). **e,** UMAP plots of two replicates of 96h Axioloids with and without MG after MNN-integration of the four samples. Note that EC-like cells only appear with MG, highlighted by black arrow head. **f,** Volcano plots in PSM and SM. Red and blue dots indicate up- and down-regulated genes by MG in both replicates, respectively. **g,** Log2 fold changes of differentially expressed genes that are up- or down-regulated by MG in SM consistently in both replicate experiments are shown. **h,** Expression levels of indicated genes in SM are compared between samples. Used abbreviations: TB (tail bud), PSM (presomitic mesoderm), aPSM (anterior presomitic mesoderm), N-SM (nascent somatic mesoderm), SM (somitic mesoderm), EC-like (endothelial cell-like), MG (Matrigel). Scale bar is 200μm.

## Matrigel only potentiates *in vitro* segmentation but does not stabilize them

Our data indicated that while MG promotes axial elongation and the initial formation of segments, it is unlikely to be sufficient to maintain or stabilize these segments. To better understand the role of MG in the elongation and segmentation of human Axioloids, we compared our scRNA-seq data of MG-containing and -lacking cultures of Axioloids. Our analysis revealed the MG-dependent emergence of an angioblast and endothelial cell-like (EC-like) population of cells at 96h of Axioloid culture (Fig.3e, Extended Data Fig.5a-d). A similar angioblast or EC-like population has been also described in trunk-like structures generated from murine pluripotent stem cells in the presence of Matrigel^3^, suggesting an evolutionary conserved role of ECM components on the genesis of vascular progenitor cells known to emerge during somitogenesis^28^.

We next further analyzed our scRNA-seq data, focusing on differentially expressed genes (DEGs) in PSM and SM cells (Fig.3f, g). Gene enrichment analysis identified up-regulation of genes associated with epithelial-to-mesenchymal transition (EMT) in SM cells of MG exposed Axioloids, while conversely PSM cells exposed to MG showed down-regulation of another distinct set of EMT associated genes (Fig.3f, g, Extended Data Fig.5e, f, Supplementary Table 2). MG-exposed SM cells furthermore showed an increase in the expression of negative regulators of WNT and TGFβ signaling pathways, *DKK1, SFPR1* and *FST, ID2* respectively (Fig.3g, h, Extended Data Fig.5e). Basement membrane associated ECM components, including *FN1* and *VIM1*, reported to be involved in the development of somitic mesoderm and known to be expressed in migratory mesenchymal cells^29–31^, were also up-regulated in SM cells of Axioloids cultured in the presence of MG (Fig.3g, h). MG exposure also changed gene expression in TB, albeit to a smaller extent than the changes in the PSM and SM of human Axioloids (Extended Data Fig.5f, g). These results, especially the identified MG-associated up-regulation of EMT signatures in SM cells, coupled with the down-regulation of epithelialization associated genes such as *TCF15^32^* and *PAX3^33^* indicate that the effect of MG on human Axioloids is not straightforward and seems to both promote and counteract molecular processes related to segmentation and epithelialization. This matches well with our observation that MG exposure leads to enhanced axial elongation and initial clear formation of segments, but then fails to stabilize these segments and is not sufficient to establish well-organized and fully epithelialized somites in Axioloids.

## RA signaling is required for segment stability and epithelialization of somites

We next looked for potential factors that may contribute to stabilizing the segmentation process and increase epithelialization of somites. Although *ALDH1A2* and molecules associated with synthesizing RA, such as *RDH10*, are specifically expressed in our system, the precursors Retinol (ROL) and Retinal (RAL) were not present in our culture conditions. Axioloids are thus unable to generate RA *de novo*, raising the question as to the function of RA signaling in Axioloids. To address this question, we added either directly RA or its precursors ROL or RAL into our *in vitro* system, during the MG embedding phase between 72h-120h. To our surprise we observed that presence of RA molecules, led to a dramatic improvement of the stability and epithelialization of forming segments within MG-embedded Axioloids at 96h and 120h. (Fig.4a, b, Extended Data Fig.6a-c, Supplementary Video 3). MG+RA Axioloids gave robustly rise to sequentially forming fully epithelialized somites with proper apical-basal polarity and central somitocoels, which MG or RA alone could not achieve (Fig.4c, Extended Data Fig.6d, e). Despite a clear effect on epithelialization, which could not be achieved by RA signaling alone (Extended Data Fig.6f), the features of these Axioloids, including overall morphology, total length, periodicity of segment formation or number of segments formed within 48h post-embedding, did not change upon addition of RA into the system (Extended Data Fig.6g-n). Assessing the effect of RA signaling on major protein and gene expression patterns within Axioloids, including that of TBXT and MEOX1, or *UNCX* and *TBX18*, we found that these molecular features were largely unchanged or even improved in RA & RAL treated Axioloids showing clear rostrocaudal identity of forming epithelial somites (Fig.4d, e, Extended Data Fig.7a-p, Supplementary Video4). MG+RA Axioloids revealed a similar change in size and expression of the TBXT^+^ SOX2^+^ TB as seen for MG only Axioloids (Extended Data Fig.7q, see also Fig.1h, Extended Data Fig.2c). Furthermore, we found that simultaneous inhibition of RA signaling via BMS493, a pan-RAR inverse agonist, while permitting RAL-mediated synthesis of RA in Axioloids, could reverse the RA effect, strongly inhibiting somite formation and epithelialization (Extended Data Fig.7r-u, Supplementary Video5).

**Fig.4.**
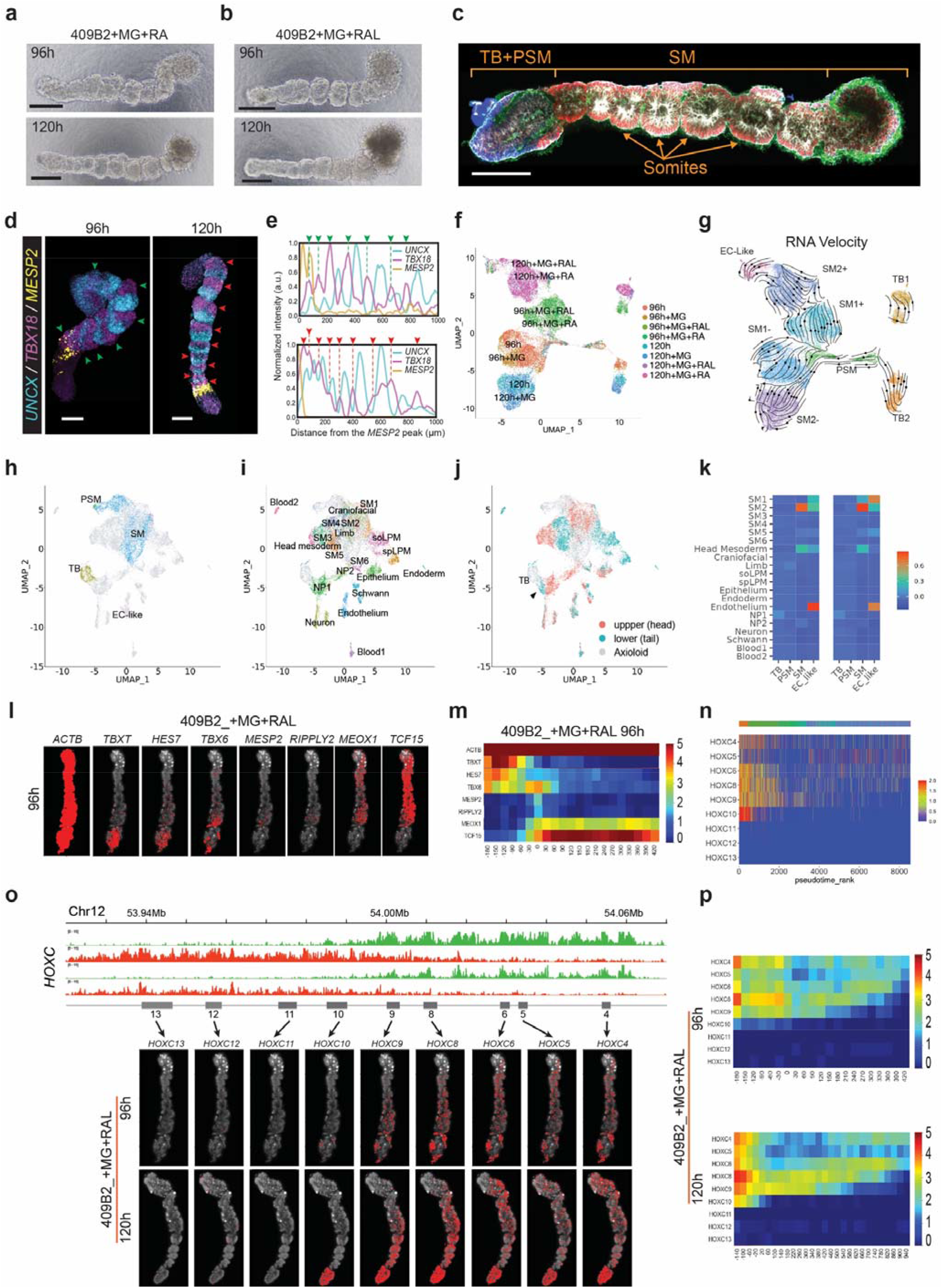
RA effect, human embryo comparison and HOX Code in Axioloids. **a** and **b,** Bright field images of Axioloids at 96h and 120h after embedding with **a,** MG and RA or **b,** MG and RAL. **c,** Immunofluorescence staining of an Axioloid embedded in MG and RA at 120h stained for F-actine (Phalloidin) in gray, FN in green, MEOX1 in red, and TBXT (BRA) in blue; shown merged channel image has been isolated from the middle of a z-stack that has been denoised using an AI based software (extracted from the **Supplementary Video 4**). **d**, and **e**, HCR staining of MG embedded Axioloids at 96h (left image) and 120h (right image) for *MESP2* in yellow, *UNCX* in cyan and *TBX18* in magenta, and corresponding signal intensity measurement along the posterior to anterior axis normalized to the position of the *MESP2* signal peak for 96h (top) and 120h (bottom). Green and red arrowheads pinpoint the *TBX18* strip pattern observed in the posterior part of each somite at 96h and 120h respectively. **f,** and **g,** UMAP projection of Axioloids at 96h and 120h, colored by samples **f,** and by clusters annotated **f.** All four conditions (MG, MG+RAL, and MG+RA) are included for each time point. **g,** RNA velocity is shown by arrows. **h-j,** UMAP projection of integrated scRNA-seq profiles of Axioloids and human embryos. Axioloids with MG and Retinal at 96h and 120h and the embryo at CS12 are analyzed. **h,** Axioloid cell clusters are colored, **i,** Cell clusters in the embryo are colored. **j,** Origins of embryonic body parts are indicated. **k,** Pearson correlation coefficient was calculated between each of the Axioloid clusters and each of the embryo clusters based on the distribution of number of cells assigned in the defined clusters for the integrated dataset. **l,** Visualization of the spatial distribution of the *ACTB, TBXT, HES7, TBX6, MESP2, RIPPLY2, MEOX1, TCF15* transcripts in a section of an Axioloid embedded in MG+RAL at 96h using HybISS and **m,** corresponding heatmap plot showing the average gene expression along the posterior to anterior axis normalized to the position of the *MESP2* signal peak (n=3). **n,** Heatmap showing the distribution of HOXC gene expression based on the pseudotime analysis of the RNA sequencing of Axioloids embedded in MG. **o,** and **p,** HOXC cluster analysis in Axioloids embedded in MG+RAL at 96h. **o,** Top panel, analysis of the epigenetic landscape at the HOXC locus profiled by CUT&Tag using antibodies against H3K4me3 (green) and H3K27me3 (red). Bottom panel, visualization of the spatial distribution of the HOXC transcripts using HybISS analysis of the HOXC cluster. **p,** Heatmap plot of the HybISS data shown in **s,** showing the average HOXC cluster gene expression along the posterior to anterior axis normalized to the position of the *MESP2* signal peak (n=3). Scale bar is 200μm.

To further dissect how RA mediates its function and alters somite formation in Axioloids, we analyzed Axioloid samples that were supplemented with RA and RAL at 96h and 120h using scRNA-seq analysis. We observed a clear segregation of the SM and TB clusters based on the presence or absence of RA and RAL, while overall identities of the cell clusters remained stable and could still be matched with each other (Fig.4f, g, Extended Data Fig.8a). Comparing the different conditions, we found that many of the DEGs, which were observed in Axioloids upon addition of MG alone, including up-regulation of EMT associated genes and of negative regulators of WNT *(SFRP1, DKK1)* or TGFβ*(FST, ID2)* signaling as well as the down-regulation of epithelialization associated transcription factors *TFC15* and *PAX3* observed in SM, were attenuated upon the addition of RA or RAL (Extended Data Fig.8b-f, Supplementary Table 3). This suggests the intriguing possibility that the presence of RA activity in the system is balancing the effect of MG, and that both, MG-mediated axial elongation and initiation of segmentation and RA-mediated stabilization and epithelialization of forming segments are required for the proper establishment and progression of somitogenesis within human Axioloids. Furthermore, we identified many known, as well as novel targets of RA signaling in the context of human segmentation and somitogenesis, including multiple RA targets in the TB and SM (Extended Data Fig.8b, c). Retinoid/RA signaling homeostasis related genes such as *RBP1, DHRS3, CYP26A1* or *CRABP2* were also upregulated by RA both in TB and SM, suggesting the presence of a negative feedback loop regulating the production and availability of RA in the system (Extended Data Fig.8e, g). Multiple transcription factors including *TCF15, MEOX1, PAX3, PBX1, ZIC3, NR2F1* and *MEIS2*, known to be expressed in somites and important for axial development of model organisms were also up-regulated in MG-embedded human Axioloids exposed to RAL/RA (Extended Data Fig.8e, h).

Our findings, showing that RA has a critical role in somite formation and epithelialization, are surprising, as the loss of RA in embryos has generally been reported to cause smaller somites or asymmetric formation of bilateral somites ^21,27,34^, rather than leading to a strong epithelialization-related phenotype. Furthermore, regarding symmetry and bilaterality in our system, we typically observed in MG and RA treated Axioloids a single axis of sequentially forming epithelial somites with single central somitocoels. Intriguingly, Axioloids also frequently displayed a superficial groove or midline-like structure, starting in the PSM and going through most of the forming segments (Extended Data Fig.9a, b), though it typically remained superficial, without separating the segments into two fully segregated somites. (Extended Data Fig.9a, b, Supplementary Video5). True bilateral, fully segregated somites, each with their respective central somitocoel, were nevertheless occasionally present in Axioloids, sometimes as an isolated fully segregated bilateral single pair of somites and in some rare cases also as a sequence of two or more somite pairs along the AP-axis of an Axioloid (Extended Data Fig.9c-e, Supplementary Video5). These bilateral somites showed usually normal protein and gene expression patterns, including proper rostrocaudal polarity (Extended Data Fig.9f-k).

## Human Axioloids mimic characteristics of human embryos

Somitogenesis is a distinctive feature of vertebrate embryos and the number of somites allows an approximate assessment of the age of an embryo. Carnegie stage (CS) 10 human embryos are characterized by 4 to 12 pairs of somites, suggesting that 96h and 120h old Axioloids are, at least, in an equivalent stage. Comparison of the dimensions of the Axioloid somites with those of CS10 & 11 embryos suggests that they are similar in shape and size (Extended Data Fig.10a-d). We furthermore found in both, Axioloids and human embryos, that earlier formed (older) somites closer to the anterior rostral region, are larger and increase in size, while newly forming somites in the caudal region have smaller volumes. The decrease in size of the posterior somites is especially evident in human Axioloids (Extended Data Fig.10a, b, Supplementary Video6). This level of high morphometric similarity observed between *in vitro-derived* Axioloids and *in vivo* human embryos, thus strikingly suggests, that even in the absence of other germ layers, the mesoderm alone can robustly self-organize and give rise to properly sized somites making up the metameric axis of the early embryo.

To further benchmark the morphogenetic and molecular features of Axioloids with that of actual human embryos, we utilized a recently uploaded scRNA-seq data set of CS12 to CS16 human embryos^35^. We reanalyzed the CS12 embryo data for comparison, and mutual nearest neighbors (MNN)-based integration analysis^36^(Extended Data Fig.10e-m) revealed, that the majority of cells present in human Axioloids matched well with the cells identified for the CS12 human embryo (Fig.4h-k, Extended Data Fig.10h-m). As expected, endodermal, neuroectodermal and other non-related mesodermal cell populations (e.g. lateral plate mesoderm (LPM), blood) identified in the human embryo, were not matching with human Axioloids (Fig.4i, k). On the other hand, Axioloids showed strong overlap with paraxial mesoderm and axial development related mesodermal cell populations found in the CS12 human embryo including populations labeled as somites or head mesoderm (Fig.4i, k, Extended Data Fig.10h-m).

We also found that a portion of Axioloid-derived angioblast/EC-like cells matched with cells marked as endothelium in the human CS12 data set (Fig.4i, k), while TB cells of Axioloids overlapped with cells labeled as neural progenitor cells (Fig.4h-k, Extended Data Fig.10e, f, h, i). As the latter population mainly matched with cells derived from the lower half of the embryo, they may represent a population similar to *SOX2/TBXT* double positive neural mesodermal progenitor cells (NMPs) found in the tail of the developing human embryo^37^ (Fig.4j).

## Presence of the HOX code in human Axioloids

Having established a robust *in vitro* system to model axial development, with the use of both MG and RA, we next looked to see whether there is a HOX Code, i.e. the spatiotemporally controlled expression of HOX genes, in human Axioloids. Combining our scRNA-seq data with HybISS-based spatial trans criptomics^38^, we successfully recapitulated the spatial expression of major TB, PSM and SM markers (Fig.4l, m) and evaluated the expression of all four HOX clusters and associated HOX genes in human Axioloids. Pseudotime analysis-based prediction of the HOX Code in Axioloids matched well with the actual spatial distribution of HOX gene expression along the anterior-posterior axis of human Axioloids (Fig.4n, Extended Data Fig.11a-c). Especially for the HOXC cluster we could observe caudal regression of HOXC genes, in line with results obtained for mouse embryos (Fig.4n-p). Combining these data sets with CUT & Tag experiments, we assessed the conducive and inhibitory epigenetic land scape of Axioloids, and observed a clear link between the switching of HOX gene expression in Axioloids and the active and repressive chromatin marks within the respective four HOX clusters, supporting the notion of a HOX Code in human Axioloids, similar to earlier findings in human and mouse gastruloids^5,39^ (Fig.4n-p, Extended Data Fig. 11d-i).

## Modulating signaling pathways in human Axioloids

Based on the morphogenetic and molecular similarities between Axioloids and actual human embryos, we then asked whether we could use the latest iteration of our model system (MG+RAL) to investigate the role of signaling pathways during human somitogenesis. Using HCR-based *in situ* hybridization, we observed that Axioloids cultured in the presence of MG+RAL, still showed clear expression gradients of *FGF8* and *WNT3a* in their TB and PSM region similar to Axioloids cultured in the presence of Matrigel only (Extended Data Fig.12a-d). RA signaling associated genes *ALDH1A2* and *CYP26A1* were also specifically expressed in their SM and TB regions respectively (Extended Data Fig.12e-h). To visualize and validate the spatial expression profiles of these and additional FGF/WNT and RA signaling pathway members predicted to exist in Axioloids from the pseudotime analysis of our scRNA-seq data, we applied HybISS-based spatial transcriptomics (Extended Data Fig.12i-l). We found that *FGF3, FGF4, FGF17, WNT5a*, and *WNT5b* showed graded expression in Axioloids similar to that of *FGF8 & WNT3a* while *RDH10* was expressed similarly to *ALDH1A2* in the SM region of MG+RAL treated Axioloids, which is consistent with previous reports in vertebrate embryos (Extended Data Fig.12i-l).

As RA signaling has a strong effect on somite formation and epithelialization, we next asked whether it also influenced the oscillation and traveling wave-like activity of the segmentation clock within Axioloids. We found that RA, RAL and ROL regardless of the presence or absence of MG in the system had largely no effect on the oscillatory activity including periodicity of the segmentation clock gene *HES7*, including robust presence of traveling wave-like expression in all treated Axioloids (Extended Data Fig.13a-j, Supplementary Video7). Inhibition of RA signaling via BMS493 had also no effect on the oscillatory activity of the segmentation clock gene *HES7* (Extended Data Fig.13k-o, Supplementary Video7). Despite the clear detrimental effect of BMS493 on epithelialization and somite formation, the overall length of BMS493 treated Axioloids remained stable (Extended Data Fig.13p-r). These findings suggest that the segmentation clock and axial elongation can be uncoupled from RA-dependent epithelial somite formation.

We then went on to modulate via small molecules also the FGF, WNT and Notch signaling pathways in human Axioloids. The observed alterations of the segmentation clock were similar to what had been previously reported *in vitro*^1,4,40^ and *in vivo*^41^. As expected, Notch inhibition with DAPT led to a quick damping and loss of oscillatory activity in Axioloids, while FGF and especially WNT inhibition had less severe effects on the segmentation clock (Extended Data Fig. 13s). The morphology of the treated Axioloids was affected and somite numbers were reduced at 120h in all three cases albeit to a smaller extent than RA inhibition (Extended Data Fig.13p, q, t). Surprisingly, FGF inhibition had the most dramatic effect on overall length of Axioloids at 120h (Extended Data Fig.13r, u, Supplementary Video8), suggesting a role of FGF signaling on axial elongation of Axioloids, similar to what has been observed in classical embryo models^42,43^.

## Modeling diseases of the human spine with Axioloids

Lastly, we investigated whether Axioloids can be used to model genetically associated diseases of the human spine. Using patient-like iPSC-lines harboring loss-of-function mutations in the coding regions of genes known to be associated with segmentation defects of the vertebrae (SDV), focusing on *HES7*^44^ or *MESP2*^45^, we generated Axioloids and assessed their morphological, molecular and functional features. We used two different *HES7* knock-out iPSC-lines which showed a similar phenotype: a conspicuous loss of segments and epithelial somite formation despite evident axial elongation (Fig.5a, b, Extended Data Fig.14a). Notwithstanding the obvious absence of segments and somites, the polarized protein expression of TBXT and MEOX1 remained largely normal (Fig.5c, Extended Data Fig. 14b, Supplementary Video9). We furthermore observed a manifest loss of rostrocaudal patterning in *HES7* KO Axioloids indicated by the loss of stripe-like expression pattern of *UNCX* and *TBX18* (Fig.5d, Extended Data Fig. 14c). We then assessed the oscillatory activity of the segmentation clock in these patient-like Axioloids lacking *HES7* and observed a clear loss of *HES7* oscillation, similar to the oscillatory phenotype, which we observed for *HES7* KO PSM cells in our previous model system^1^ (Fig.5e-g, Extended Data Fig.14d, Supplementary Video 10).

**Fig.5.**
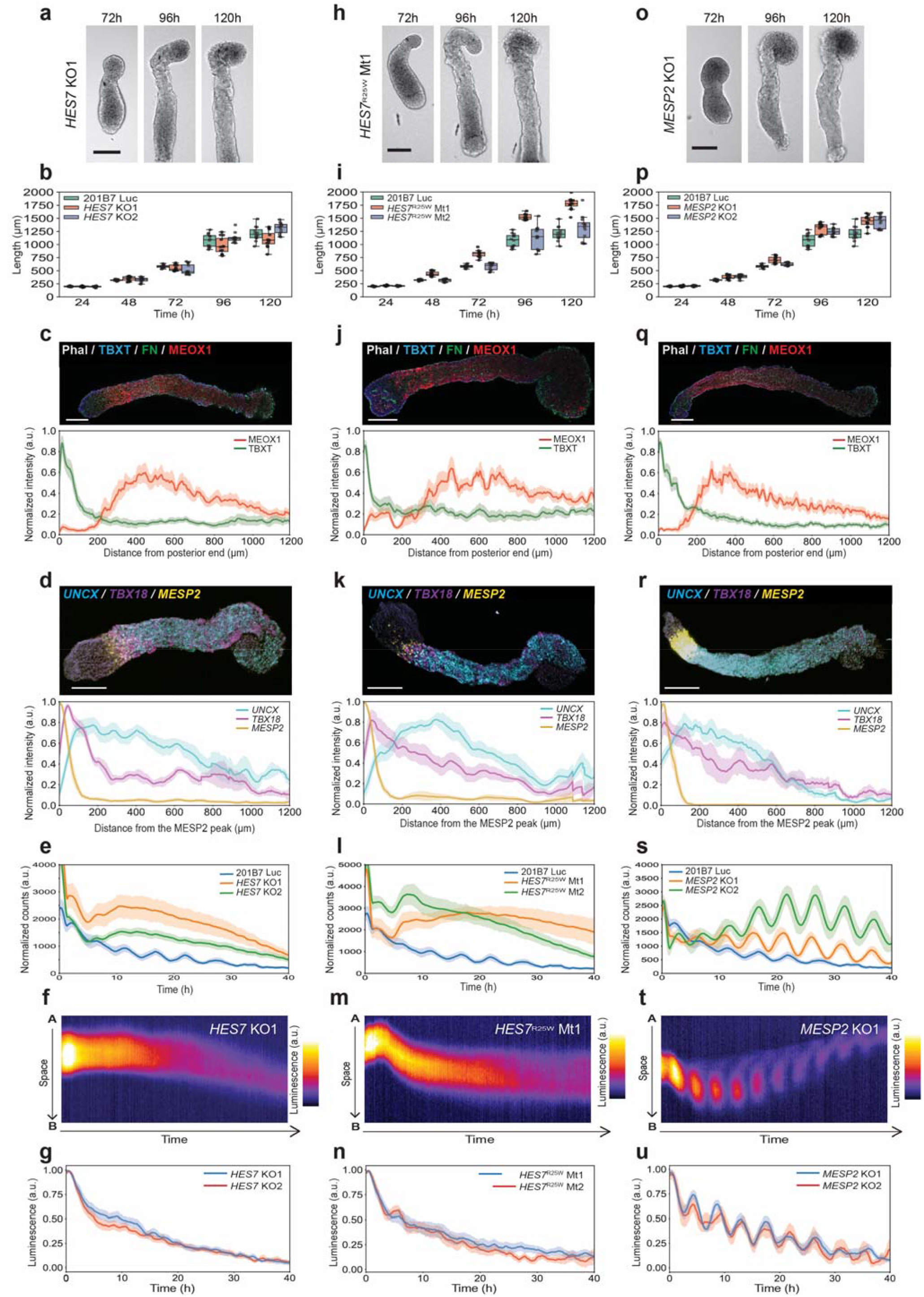
Molecular and functional characterization of patient-like Axioloids. Panels **a-g,** show data for *HES7* KO1, panels **h,** to **n,** show data for *HES7*^R25W^ MT1 and panels **o-u,** show data for *MESP2* KO1. **a, h,** and **o,** Serial brightfield images of a forming patient-like Axioloid at 72h, 96h and 120h extracted from **Supplementary Video 9, 11,** and **13** respectively. **b, i,** and **p,** Axioloid length along the posterior to anterior axis at 24h, 48h, 72h, and at 96h and 120h for the corresponding cell lines *HES7* KO1 and *HES7* KO2 (N=3, n=12), *HES7*^R25W^ MT1 and *HES7*^R25W^ MT2 (N=3, n=12), *MESP2* KO1 and *MESP2* KO2 (N=3, n=12). **c**, **j,** and **q,** Immunofluorescence staining and signal quantification of MG+RAL embedded Axioloids at 120h. Top panel, merged channel images of Axioloids stained for F-actine (Phalloidin) in gray, FN in green, and MEOX1 in red, TBXT (BRA) in Blue. Bottom panel, corresponding quantification along the posterior to anterior axis of TBXT (green line) and MEOX1 (in red) signal intensity for *HES7* KO1 (N=3, n=10), *HES7^RyW^* MT1 (N=4, n=9), *MESP2* KO1 (N=3, n=12). **d, k,** and **r,** HCR staining and signal quantification of MG+RAL embedded Axioloids at 120h. Top panel, *MESP2* in yellow, *UNCX* in cyan and *TBX18* in magenta, and bottom panel, corresponding signal intensity measurements along the posterior to anterior axis normalized to the position of the *MESP2* signal peak for *HES7* KO1 (N=3, n=9), *HES7^R25W^* MT1 (N=3, n=8), *MESP2* KO1 (N=3, n=7). **e, l,** and **s,** Time series of the HES7:Luciferase expressing hiPSC cell line (209B7 Luc) from 72h to 112h for *HES7* KO1 (N=4, n=12), *HES7* KO2 (N=4, n=12), *HES7*^R25W^ MT1 (N=3, n=9), *HES7^R25W^* MT2 (N=3, n=9), *MESP2* KO1 (N=4, n=12), *MESP2* KO2 (N=4, n=12). **f, m,** and **t,** Kymographs of **f,***HES7* KO1, **m,***HES7*^R25W^ MT1, and **t,***MESP2* KO1 Axioloids embedded in MG+RAL. **g, n,** and **u,** Average measured HES7:Luciferase signal over time for *HES7* KO1 (N=5, n=8), *HES7* KO2 (N=6, n=9), *HES7*^R25W^ MT1 (N=5, n=10), *HES7*^R25W^ MT2 (N=5, n=9), *MESP2* KO1 (N=3, n=5), *MESP2* KO2 (N=3, n=7). Scale bar is 200μm.

We observed a similar range of phenotypes in Axioloids derived from patient-like iPSC lines containing a point mutation (rs113994160: c.73C>T) in *HES7*, resulting in a pathogenic missense mutation R25W in the helix-loop-helix domain of HES7 reported to cause segmentation defects of the vertebrae (SDV)^44^ (Fig.5h-n, Extended Data Fig.14e-h, Supplementary Videos11 & 12). Axial elongation in these *HES7^R25W^* Axioloids appeared to be slightly increased as compared to the healthy donor control iPSC-derived Axioloids (Fig.5h, i, Extended Data Fig.14e). *HES7^R25W^* MT1 & MT2 derived *HES7*^R25W^ exhibited normal expression of TB & SM markers TBXT and MEOX1 (Fig.5j, Extended Data Fig.14f, Supplementary Video 11) combined with clear loss of rostrocaudal patterning (Fig.5k, Extended Data Fig.14g), and loss of *HES7*oscillation, again matching our findings for *HES7* KO Axioloids as well as our previously reported oscillatory phenotype for *HES7^R25W^* iPSC-derived PSM cells^1^ (Fig.5l-n, Extended Data Fig.14h, Supplementary Video 12).

Next, we assessed the effects of the loss of *MESP2*, an aPSM associated transcription factor for which pathogenic mutations in patients with SDV have been previously reported^45^. Using *MESP2* knock-out iPSC lines *(MESP2* KO1 & *MESP2* KO2) we derived patient-like Axioloids and again assessed various morphological, molecular and functional features. *MESP2* KO Axioloids elongated normally but were devoid of segments or epithelial somites (Fig.5o, p, Extended Data Fig.14i). In contrast to the largely normal expression pattern of TBXT and MEOX1, *MESP2* KO Axioloids (Fig.5q, Extended Data Fig.14j, Supplementary Video 13) exhibited severely impaired rostrocaudal patterning, characterized by the lack of stripe-like expression of *UNCX* coupled with low-level expression of *TBX18* (Fig.5r, Extended Data Fig.14k). This *in vitro*“human phenotype” resembled strikingly the one reported for *MESP2* KO mice^46,47^, indicating that Axioloids can reconstitute complex genetic phenotypes *in vitro*. In contrast to *HES7*, loss of *MESP2* did not lead to a loss of *HES7* oscillatory activity in Axioloids (Fig.5s-u, Extended Data Fig.14l, Supplementary Video 14). Together, this data shows that Axioloids can provide not only invaluable insights into normal human axial development but also contribute to a better understanding of the pathogenesis of diseases of the human spine.

In summary, we have established and characterized in-depth a pluripotent stem cell-derived 3D mesodermal model of human axial development, which could reconstitute various aspects of human somitogenesis and axial development *in vitro*. Axioloids recapitulated a range of complex developmental processes including axial elongation, segmentation, epithelialization to oscillation of the segmentation clock, while also sharing molecular and morphometric features with the tail and axis of the developing human embryo. Our bottom-up approach revealed the remarkable self-organization potential of primitive and paraxial mesoderm, which can give rise to the metameric basic body plan of the human embryo even in the absence of other germ layers. We also uncovered a crucial role of RA signaling on the morphogenetic processes associated with segmentation and somite formation within Axioloids, suggesting that RA supplementation especially in combination with ECM components might also improve the morphogenetic features of other *in vitro* model systems of human and non-human early embryonic development.

Our bottom-up experimental approach demonstrates that complex developmental events such as somitogenesis, can be deconstructed and dissected into discrete “building blocks” of developmental principles which are usually intricately connected and cannot be easily uncoupled *in vivo*. Axioloids, a self-organizing *in vitro* model of human axial development allowed us to individually assess and manipulate such building blocks at the molecular, cellular and morphogenetic level. Further iterations of this model system will likely incorporate still “missing” anatomical structures such as notochord or neutral tube, which will allow assessment of subsequent stages of somitic development and differentiation including compartmentalization of somites into sclerotome, dermomyotome and other functional derivatives. Axioloids, a surrogate model of the human embryonic tail and forming axis, are capable of recapitulating core features of human somitogenesis, and represent an exciting new platform to investigate axial development and disease in a human context.

## Data availability

All single cell RNA sequencing data and CUT&Tag data used for this study have been deposited in the NCBI Gene Expression Omnibus under accession number GSE199576.

## Code availability

Computational codes and scripts used in this study are available at GitHub (https://github.com/Alev-Lab/Axioloids-manuscript.git) and upon request from the corresponding author.

## Acknowledgements

The authors thank G. Sheng and S. Goulas for critical reading of the manuscript; members of the single cell core facility (SignAC) at ASHBi and S. Terakura for help with scRNA-seq experiments; K. Terai and other members of the Matsuda lab at Kyoto University for help with bioluminescence live imaging experiments; members of the iCeMS Analysis Center and Y. Tomoda and S. Kihara from the CiRA imaging core facility for support with microscopy and high content live imaging; Y. Arai and members of the ASHBi office for administrative support. This work was supported by Naito Foundation Scientific Research Grant to C.A.; Takeda Science Foundation Grant to C.A.; Japan Agency for Medical Research and Development (AMED) Grant Number JP20bm0704050 to C.A.; the ISHIZUE 2020 of Kyoto University Research Development Program to C.A.; JST FOREST Program (Grant Number JPMJFR206C, Japan) to T.Y.; JST, CREST (Grant Number JPMJCR2023, Japan) to T.Y.; the ASHBi Fusion Research Program to K.YK., Y.Y., K.S., T.Y., C.A.; N.M is supported by the Francis Crick Institute which receives its core funding from Cancer Research UK (FC011181), the UK Medical Research Council (FC011181), and the Wellcome Trust (FC011181); a Royal Society research grant (RGS\R2\212082); and A.MA., N.M. and C.A. where additionally supported by the UK Medical Research Council and Japanese Agency for Medical Research and Development AMED (MR/V005367/2). ASHBi is supported by the World Premier International Research Center Initiative (WPI), MEXT, Japan.

## Author contributions

Y.Y., K.YK., S.M., S.H. and C.A. developed human Axioloid induction protocol and performed molecular and functional analysis of samples; A.MA. and N.M provided critical scientific feedback and frequent discussions; Y.Z. helped with molecular characterization of samples and data analysis; K.S. performed HybISS analysis of samples with the help of Y.Y. and S.H.; J.L.T. repeated Axioloid experiments in the UK, with support from N.M.; J.K. and S.L. provided data of CS10 & CS11 human embryo samples; S.H. quantified somite dimensions in Axioloids and human embryos; S.K. supported microscopy and live imaging of samples; A.M., Y.K. and T.T. performed and analysed CUT&Tag experiments; Y.Y. performed and T.T., Y.K. and T.Y. analysed scRNA-seq data; C.A. conceived, designed and supervised the study; C.A. analysed and interpreted the data and wrote the manuscript incorporating feedback from co-authors. All authors discussed and commented on the manuscript and agreed on the presented results.

---

**Extended Data Fig.1.**
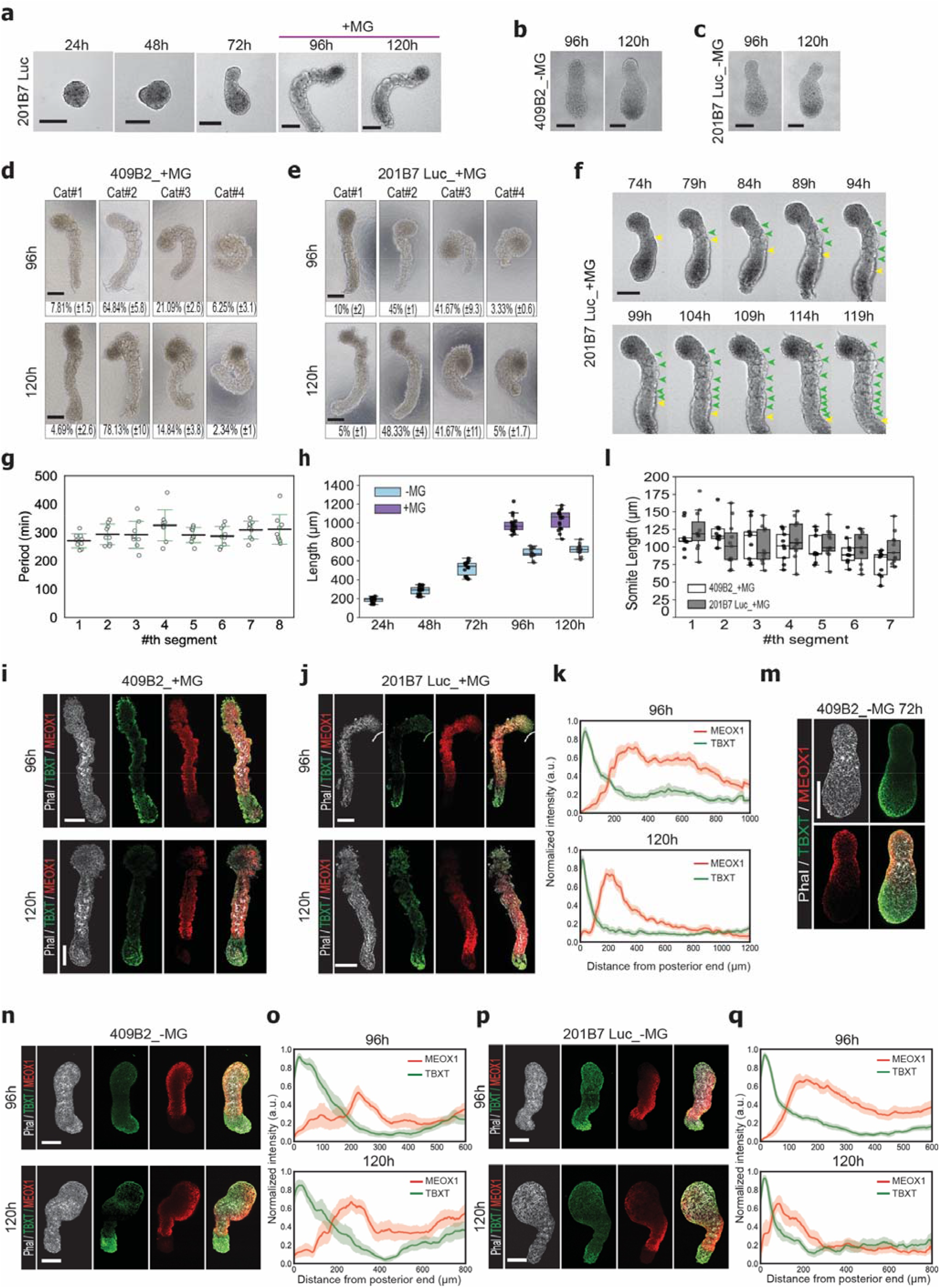
Morphological and molecular characterization of human Axioloids. **a-f,** Bright field images of **a,** elongating Axioloids at 24h, 48h, and 72h, followed by images of Axioloids after MG embedding at 96h and 120h or **b, c,** without MG embedding. **d,** and **e,** Representative bright field images reflecting the different morphology-based categories (Cat#1: straight, minimal curvature, clear segments; Cat#2: curved but clear segments; Cat#3: very curved, segment borders are still distinguishable; Cat#4: not properly elongated or completely collapsed; segment borders not clear) of MG embedded Axioloids at 96h and 120h in 2 different cell lines **d,**409B2 (N=3, n=128 Axioloids) and **e,**201B7 Luc (N=3, n=60 Axioloids). **f**, Serial images of a forming Axioloid at 5h intervals, from 74h to 119h (extracted from **Supplementary Video 1**). Colored arrowheads highlight the process of segment formation at each shown time point with the yellow arrowheads pinpointing areas where somite segmentation is ongoing whereas green arrowheads highlight the areas where segmentation is completed. **g,** Periodicity of somite segmentation based on live-cell imaging observations (N=3, n=9). **h,** Axioloid length (antero-posterior) at 24h, 48h, 72h, and at 96h and 120h with or without MG embedding (24h, 48h, 72h, N=3, n=18, and =MG 96h n=14, +MG 96h n=18, and -MG 120h n=13, +MG 120h n=18). **i-q,** Immunofluorescence staining and corresponding quantifications. **i, j, m, n, p,** Representative images of F-actine (Phalloidin) in gray, TBXT in green, and MEOX1 in red stained Axioloids at **m,** 72h, **i, j,** 96h and 120h with MG embedding and **n, p,** 96h and 120h without MG embedding. **k, o, p,** Corresponding quantification along the posterior to anterior axis of TBXT (green line) and MEOX1 (in red) signal intensity at **k,** 96h (N=3, n=14) and 120h (N=4, n=16) with MG embedding and **o, q,** 96h (409B2 N=3, n=9 and 201B7 N=3, n=15) and 120h (409B2 N=3, n=9 and 201B7 N=3, n=11) without MG embedding. **l,** Somite length was measured based on Phalloidin staining done in **i,** and **j,** in both cell lines 409B2 (N=3, n=9) and 201B7 Luc (N=3, n=9). Scale bar is 200μm.

**Extended Data Fig.2.**
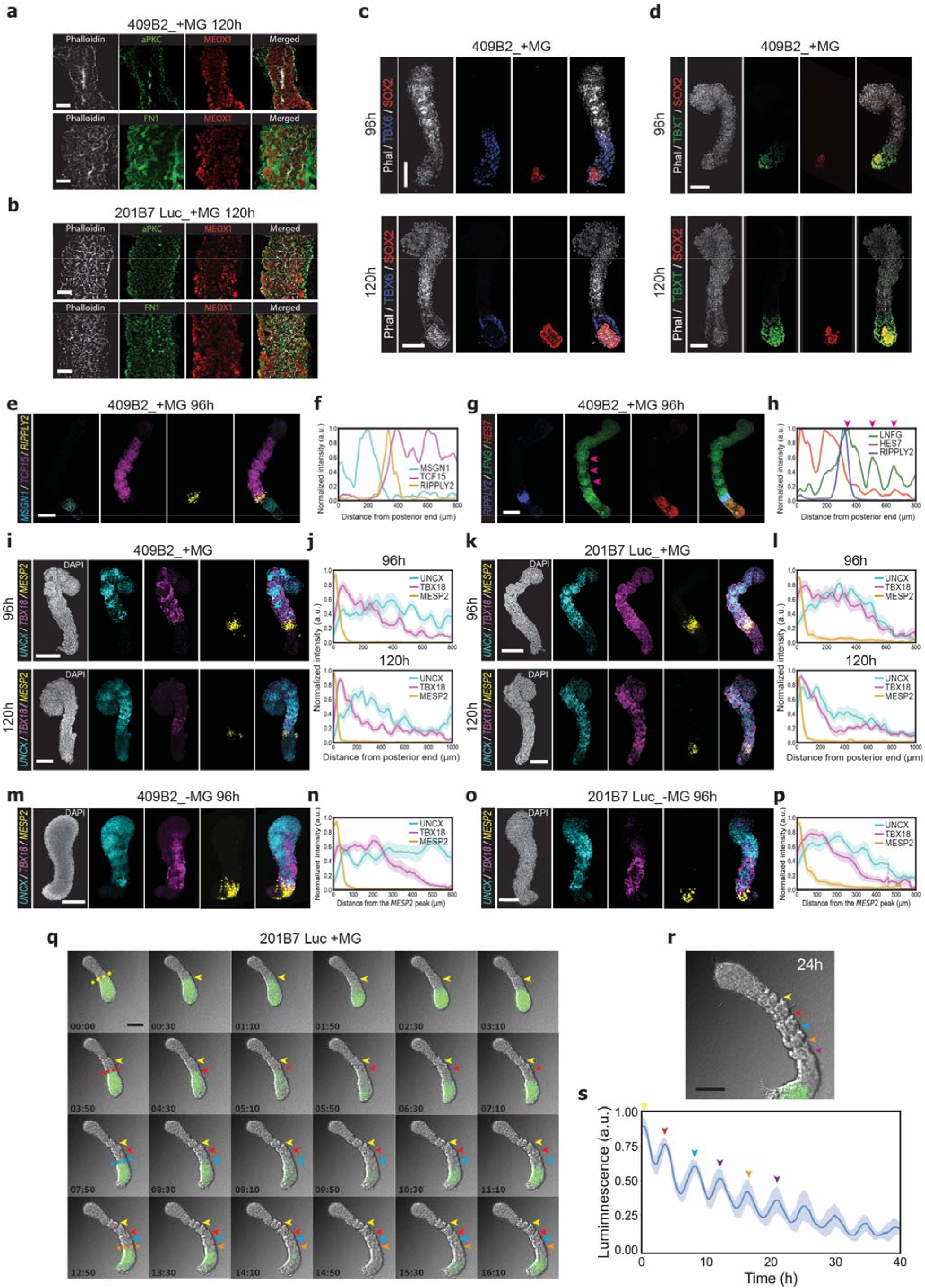
Assessment of apicobasal polarity, developmental protein-& gene expression patterns, rostrocaudal patterning, traveling wave front of *HES7* oscillatory activity & segmentation in human Axioloids embedded in MG. **a-d,** Immunofluorescence staining of MG embedded Axioloids. **a,** and **b,** High magnification images (X63) of a single segment at 120h with F-actine (Phalloidin) in gray, aPKC and FN in green, and MEOX1 in red in **a,** 409B2 and **b,** 201B7 Luc iPS cell line-derived Axioloids; images are representative of 2 independent experiments. **c,** Images of F-actine (Phalloidin) in gray, TBX6 in blue and SOX2 in red and **d,** Images of F-actine (Phalloidin) in gray, TBXT in green and SOX2 in red. **e-p,** HCR staining and corresponding signal quantification of **e,** and **f,** HCR staining of *MSGN1* in cyan, *TCF15* in magenta and *RIPPLY2* in yellow and **g,** and **h,** of *RIPPLY2* in blue, *LFNG* in green and *HES7* in red; the purple arrowhead highlights the stripe-like staining pattern of *LFNG* in the posterior half of each somite, shown images are representative of 2 independent experiments. **i-p,** HCR staining of *MESP2* in yellow, *UNCX* in cyan and *TBX18* in magenta, and corresponding signal intensity measurements along the posterior to anterior axis normalized to the position of the *MESP2* signal peak of Axioloids embedded in MG at 96h and 120h respectively in 2 different cell lines **i**, **j**, 409B2 (N=4, n=10 and N=3, n=9) and **k, l,**201B7 Luc (N=3, n=9 and N=3, n=6) and Axioloids without MG embedding at 96h in 2 cell lines **m**, **n**, 409B2 (N=3, n=12) and **o, p,** 201B7 Luc (N=3, n=11). Single channel images shown in **c-e, g,** and **i,** top panel, corresponds to the merged channel images shown in Fig.1 **h-j**. **q,** and **r,** Annotated serial images of a forming Axioloid with HES7:Luciferase signal overlayed in green (extracted from **Supplementary Video 2**). Colored dotted lines mark the furthermost anterior position reached by each HES7 oscillation wave of gene expression, identical colored arrowheads mark this position overtime, yellow first oscillation, red second, blue third, orange forth. **r,** Image at 24h shows that each HES7:Luciferase expression wavefront position corresponds to the area of a formed segment. **s,** Average HES7:Luciferase intensity measurements overtime (N=4, n=8). Scale bars in **a,** and **b,** 50μm others are 200μm.

**Extended Data Fig.3.**
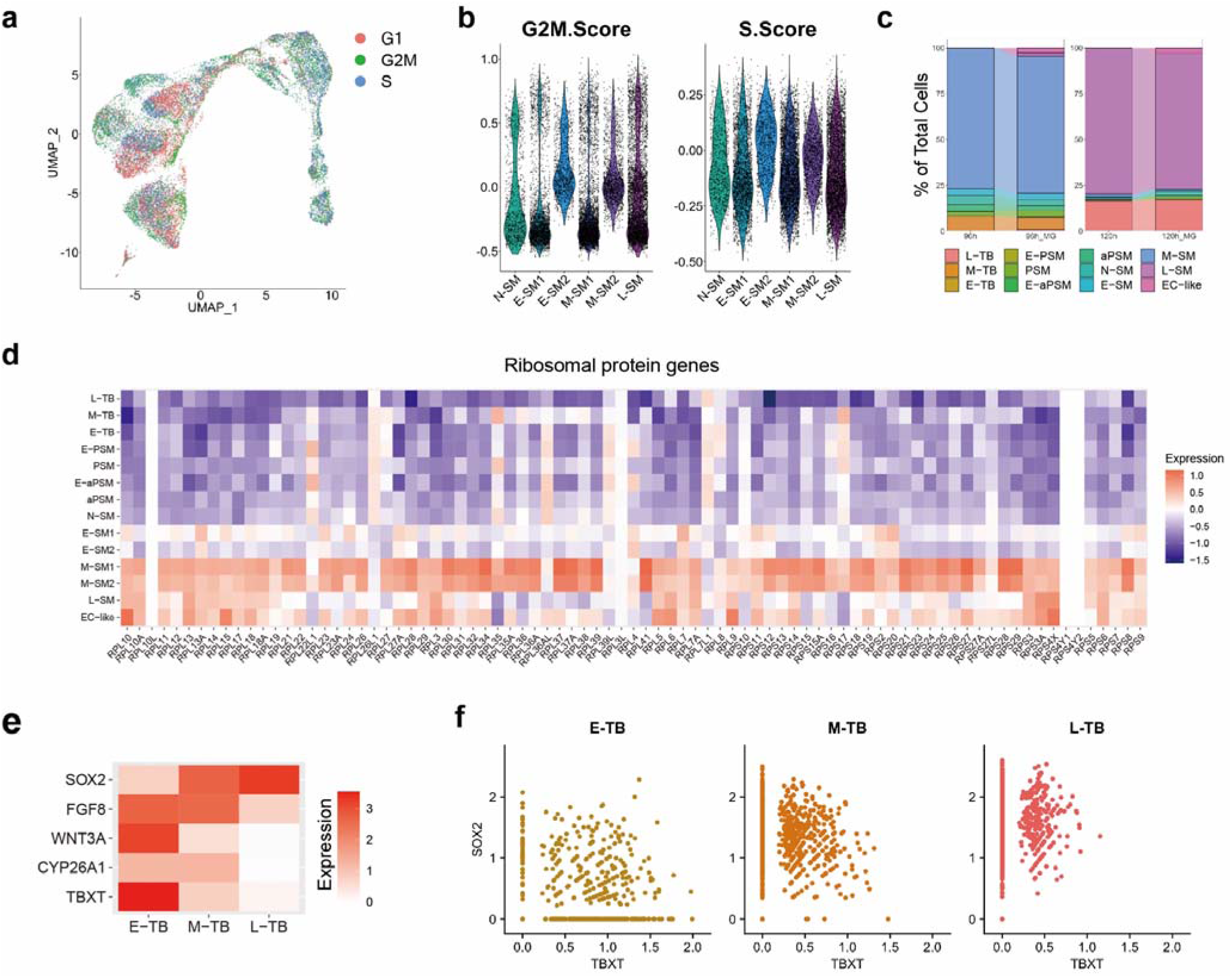
Single cell RNA-seq analysis of human Axioloids. **a,** UMAP projection of scRNA-seq datasets of Axioloids at 48h, 72h, 96h and 120h, colored by inferred cell cycle phases (G1, G2M, S). **b,** G2M.Score and S.Score of the cells in each cluster of Fig.2b. **c,** Proportions of cell types in Axioloids with and without MG for both 96h and 120h timepoints. **d,** Averaged expression levels of ribosomal protein genes in each cluster of Fig.2b. **e,** Transition of TB marker gene expression along the time course. **f,** Expression levels of TBXT and SOX2 in each cell are plotted for the three TB clusters (E-TB, M-TB and L-TB, standing for early, mid and late tailbud respectively).

**Extended Data Fig.4.**
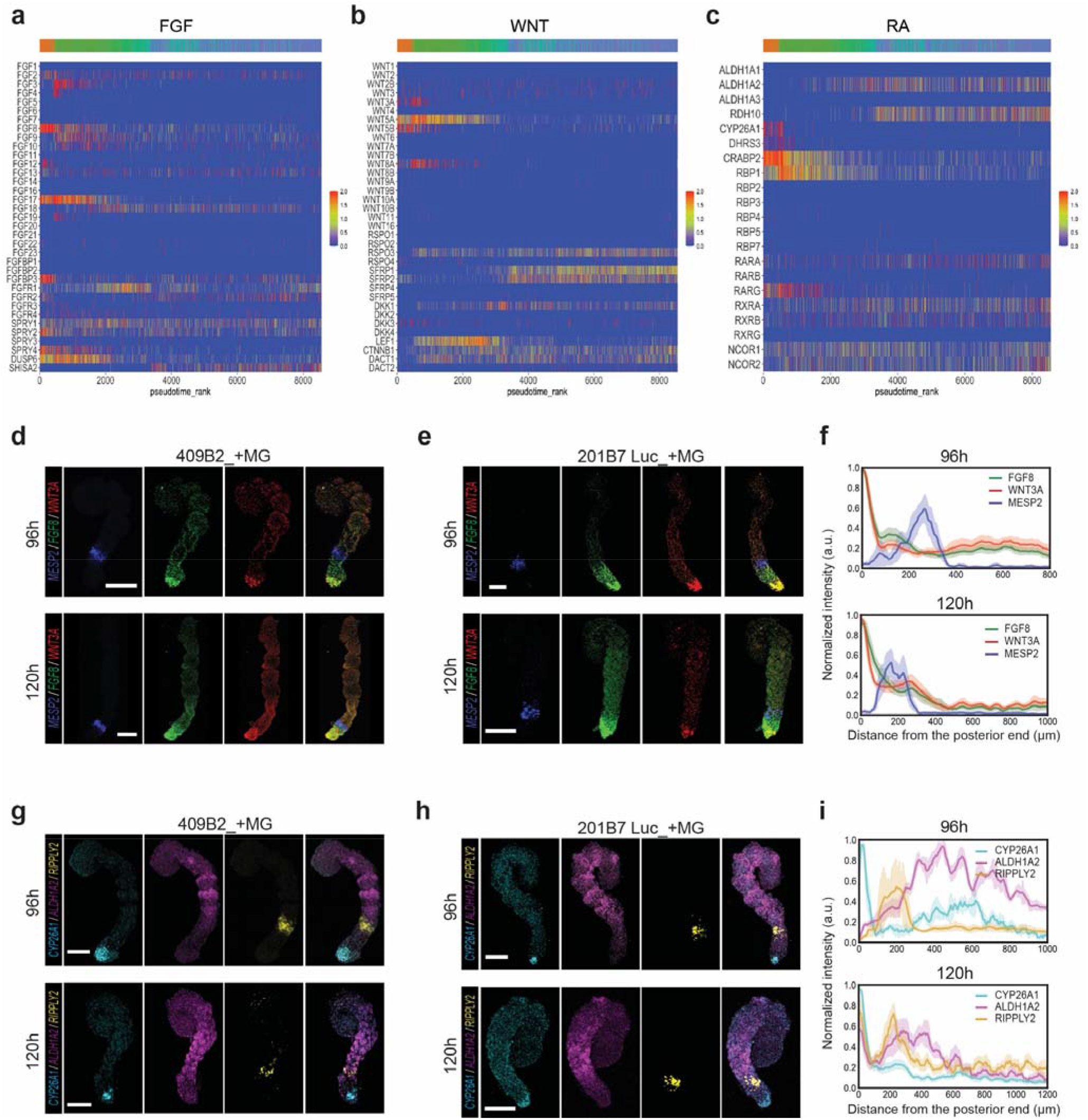
Expression gradients of FGF, WNT and RA signaling pathway members in human Axioloids embedded in MG. **a-c,** Pseudotime representation of expression of FGF, WNT and RA signaling pathway associated transcripts in human MG exposed Axioloids at 96h of culture (24h after embedding into MG); gene expression patterns arranged along pseudotime rank. Pseudotime expression patterns for effectors and negative regulators of all three pathways are included. **d-i,** HCR staining images and signal quantification of MG embedded Axioloids at 96h and 120h derived from **d,** and **g,** 409B2 and **e, f, h,** and **i,** 201B7 Luc iPSC lines. Shown images are representative of 3 independent experiments. **d,** and **e,** HCR staining of *MESP2* in blue, *FGF8* in green and *WNT3a* in red. **f,** corresponding quantification along the posterior to anterior axis of the signal intensity normalized the position of the *WNT3a* signal peak (96h: N=3, n=8 and 120h: N=3, n=7). **g,** and **h,** HCR staining of *CYP26A1* in cyan, *ALDH1A2* in magenta and *RIPPLY2* in yellow and **i,** corresponding quantification along the posterior to anterior axis of the signal intensity normalized the position of the *CYP26A1* signal peak (96h: N=3, n=6 and 120h: N=3, n=7). Scale bar is 200 μm.

**Extended Data Fig.5.**
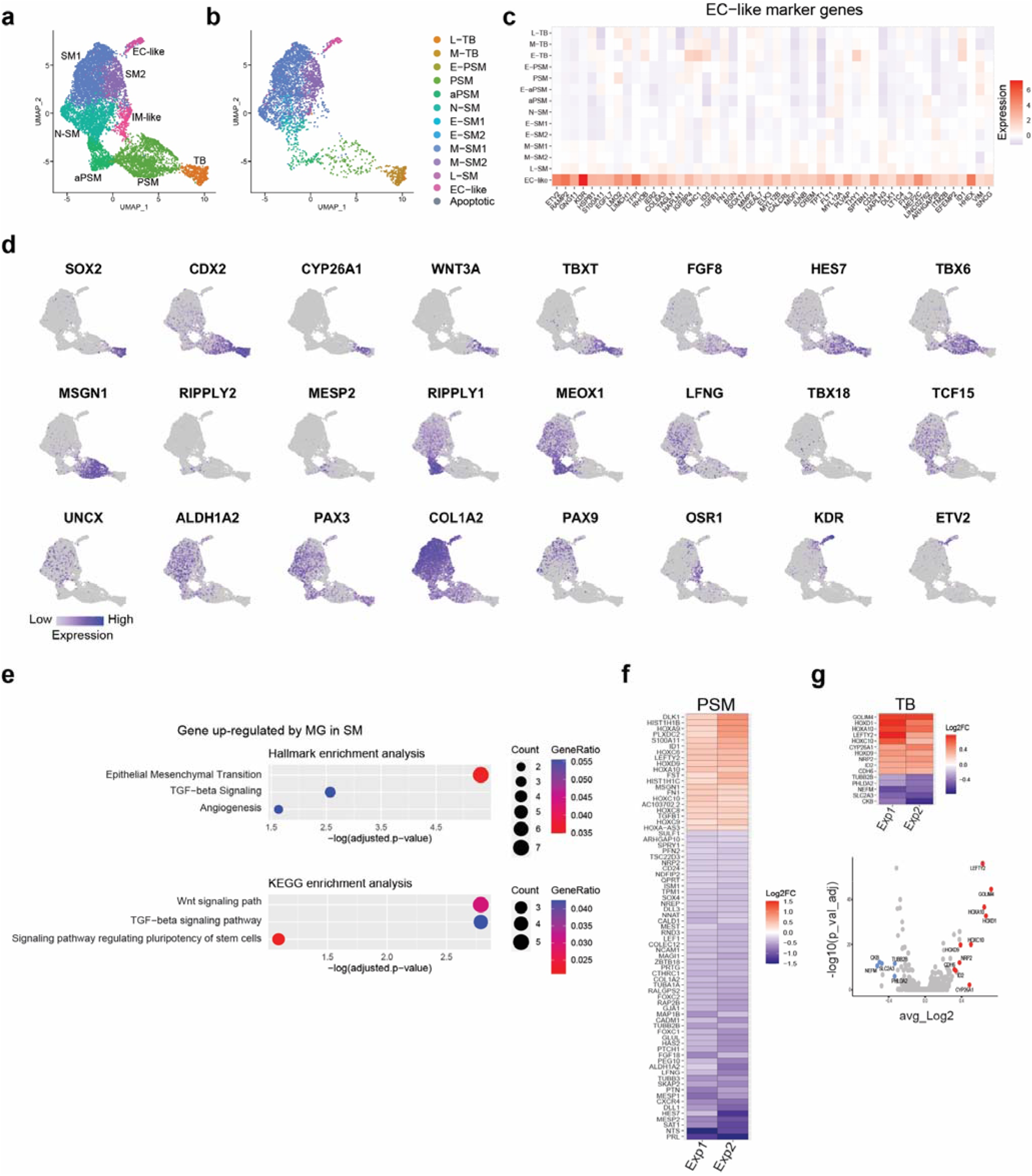
Single cell RNA-seq analysis: identification of DEGs associated with the exposure of Axioloids to MG. **a,** UMAP projection of the integrated two replicates of Axioloids at 96h with MG, colored by the clusters of Fig.2g. **b,** UMAP projection of the integrated two replicates of Axioloids at 96h with MG, colored by the clusters of Fig.2b. Note that **b,** includes only replicate 1. **c**, Expression level of indicated genes on the same UMAP plot in **a**. **d,** Averaged expression levels of identified EC-like marker genes in each cluster of Fig.2b. **e,** Enrichment analysis of upregulated genes in SM in the presence of MG. Hallmark gene sets and KEGG datasets are used. **f,** and **g,** Differentially expressed genes between Axioloids with and without MG at 96h in PSM and TB.

**Extended Data Fig.6.**
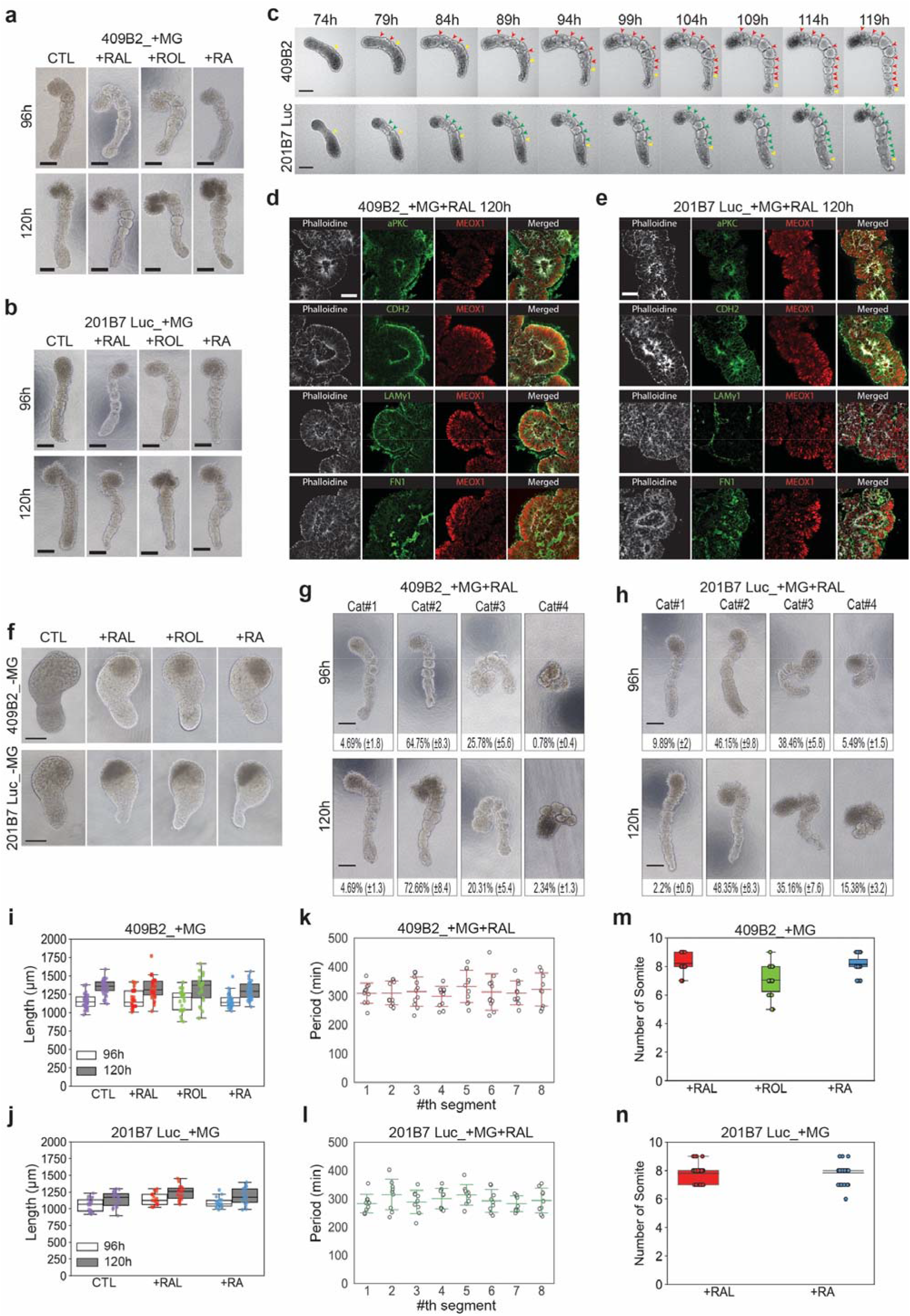
Assessing the morphogenetic effects of retinoid signaling on human Axioloids. **a,** and **b,** Bright field images of Axioloids at 96h and 120h after embedding in MG (Matrigel) only, MG+RAL (Retinal), MG+ROL (Retinol) or MG+RA (Retinoic Acid) for **a,**409B2 and **b,** 201B7 Luc. **c,** Serial images of an elongating Axioloid at 5h intervals, from 74h to 119h (extracted from **Supplementary Video 3**). Colored arrowheads highlight the process of segment formation at each shown time point with the yellow arrowheads pinpointing to areas where somite segmentation is ongoing whereas red for 409B2 or green for 201B7 Luc arrowheads highlight the areas where segmentation is completed. **d,** and **e,** Immunofluorescence high magnification images (X63) of a single somite of Axioloids embedded in MG+RAL at 120h with F-actine (Phalloidin) in gray, and from top to bottom aPKC, CDH2, LAMC1, FN in green, and MEOX1 in red in **d,** 409B2 and **e,** 201B7 Luc iPS cell lines; images are representative of 2 different experiments. **f,** Bright field images of Axioloids at 120h without MG embedding, after addition of RAL only, ROL only and RA only (no MG in all cases) in 409B2, top panel and 201B7 Luc, bottom panel. **g,** and **h,** Representative bright field images reflecting the different morphology-based categories (Cat#1: straight, minimal curvature, clear somites; Cat#2: curved but clear segments & somites; Cat#3: very curved, somites & segment borders are still distinguishable; Cat#4: not properly elongated or completely collapsed; somites & segment borders not clear) of MG+RAL embedded Axioloids at 96h and 120h in 2 different cell lines **g,** the 409B2 (N=5, n=128 Axioloids) and **h,** the 201B7 Luc (96h N=3, n=91 Axioloids and 120h N=3, n=92 Axioloids). **i,** and **j,** Length measurement of Axioloids at 96h and 120h after embedding in MG only, MG+RAL, MG+ROL or MG+RA for **i,** 409B2 (N=3, +MG 96h n=28, 120h n=31, +MG+RAL 96h n=27, 120h n= 30, +MG+ROL 96h n=21, 120h n=20, +MG+RA 96h n=20, 120h n=120h) and **j,** 201B7 Luc (N=3, +MG 96h n=18 otherwise n=17). **k,** and **l,** Periodicity of somite segmentation based on live-cell imaging observations **k,** for 409B2 (N=3, n=11) and **l,** 201B7 Luc (N=3, n=9) respectively. **m,** and **n,** Total number of somites in Axioloids embedded in **m,** MG+RAL or MG+ROL or MG+RA at 120h in 409B2 (N=3, n=39/39 and N=2, n=25 respectively) and **n,** MG+RAL or MG+RA at 120h in 201B7 Luc (N=3, n=60 for RAL and n=59 for RA). Scale bar in **d,** and **e,** is 50μm, others are 200μm.

**Extended Data Fig.7.**
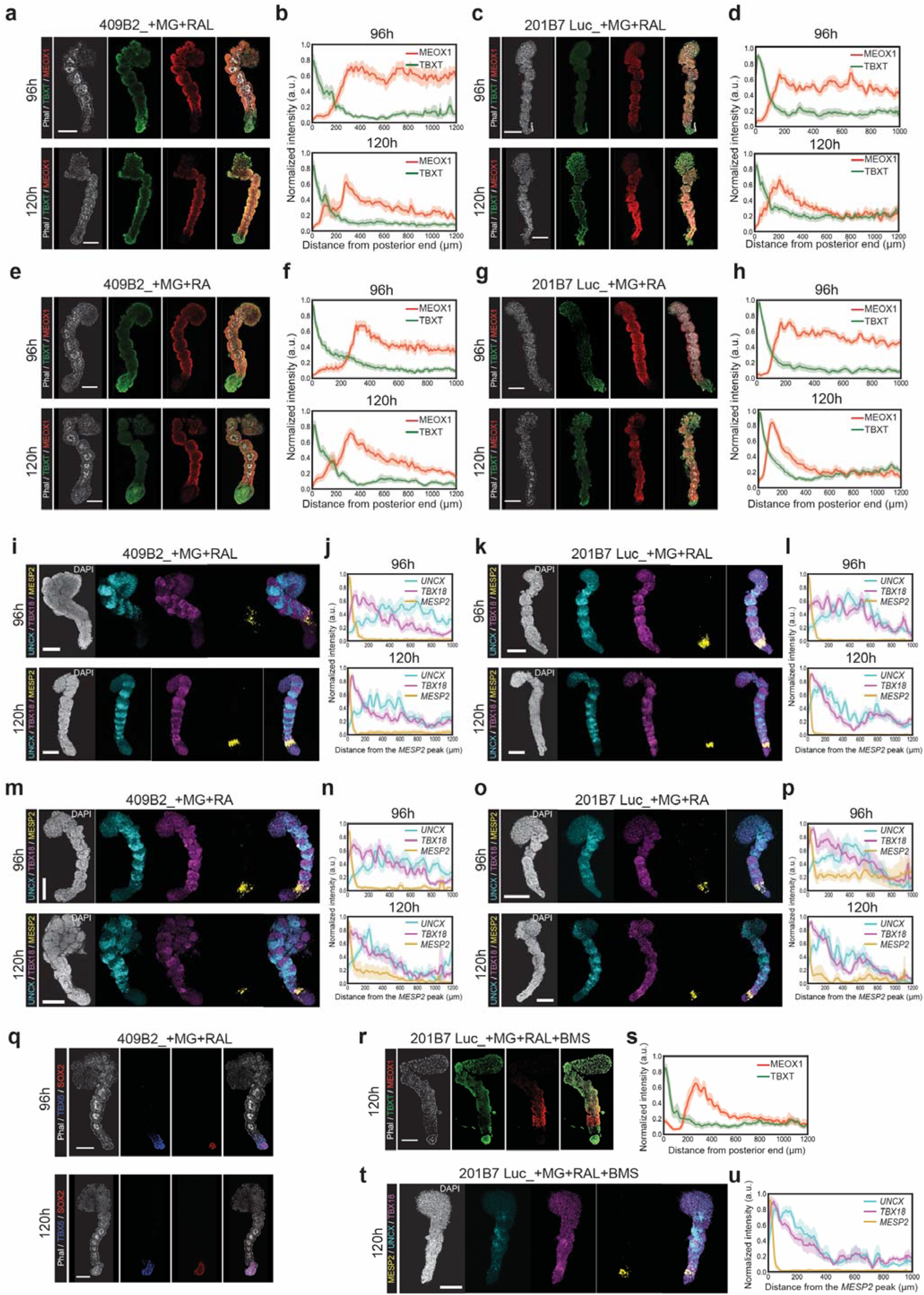
Molecular characterization of human Axioloids exposed to agonists or inhibitors of Retinoic Acid (RA) signaling. **a**-**h**, Representative images of immunofluorescence staining of F-actine (Phalloidin) in gray, TBXT in green, and MEOX1 in red and corresponding quantification of signal intensity of Axioloids embedded in MG+RAL **a**, **b**, 409B2 at 96h (N=4, n=10) and 120h (N=4, n=9) and **c**, **d**, 201B7 Luc at 96h (N=4, n=13) and 120h (N=4, n=13). Immunofluorescence data of Axioloids embedded in MG+RA shown in **e**, **f**, 409B2 at 96h (N=4, n=15) and 120h (N=4, n=12) and **g**, **h**, 201B7 Luc at 96h (N=3, n=15) and 120h (N=4, n=16). **i**-**p**, Representative images of HCR staining of *MESP2* in yellow, *UNCX* in cyan and *TBX18* in magenta, and corresponding signal intensity measurement along the posterior to anterior axis normalized to the position of the *MESP2* signal peak of Axioloids embedded in MG+RAL **i**, **j**, 409B2 at 96h (N=3, n=9) and 120h (N=3, n=9) and **k**, **l**, 201B7 Luc at 96h (N=3, n=9) and 120h (N=3, n=9). In situ hybridization data of Axioloids embedded in MG+RA **m**, **n**, 409B2 at 96h (N=3, n=9) and 120h (N=4, n=8) and **o**, **p**, 201B7 Luc at 96h (N=2, n=5) and 120h (N=2, n=5). **q,** Immunofluorescence staining of F-actine (Phalloidin) in gray, TBXT in blue, and SOX2 in red of 96h (top), and 120h (bottom), 409B2 Axioloids embedded in MG+RAL. Images shown are representative of 3 independent experiments. Scale bar is 200μm. **r,** and **s,** Representative images of immunofluorescence staining of F-actine (Phalloidin) in gray, TBXT in green, and MEOX1 in red and corresponding quantification of signal intensity of Axioloids embedded in MG+RAL+BMS493 (201B7 Luc) at 120h of culture (N=3, n=11). **t**, and **u**, Representative images of HCR staining of *MESP2* in yellow, *UNCX* in cyan and *TBX18* in magenta, and corresponding signal intensity measurement along the posterior to anterior axis normalized to the position of the *MESP2* signal peak of Axioloids embedded in MG+RAL+BMS493 (201B7 Luc) at 120h of culture (N=2, n=6).

**Extended Data Fig.8.**
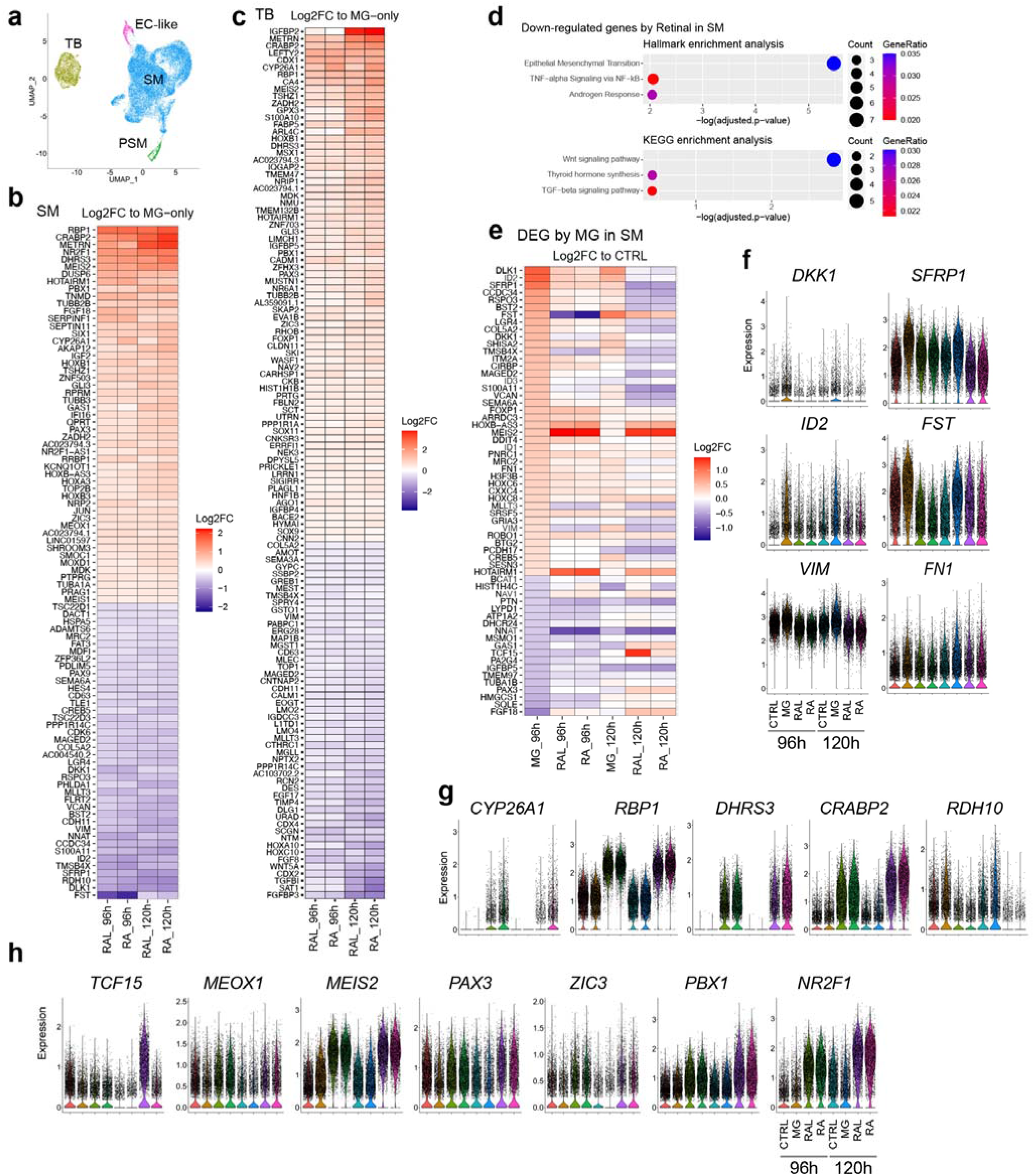
Single cell RNA-seq based assessment of RA signaling effect on MG embedded human Axioloids. **a,** Integrated UMAP projection of single-cell transcriptome profiles of Axioloids with all the four conditions (control, MG only, MG+RAL (Retinal) and MG+RA (Retinoic Acid) at both 96h and 120h. **b,** and **c,** Expression changes by MG+RAL (Retinal) and MG+RA (Retinoic Acid) compared to the MG only conditions for consistently up- or down-regulated genes at both 96h and 120h in SM **b,** and TB **c**. **d,** Enrichment analysis of down-regulated genes in SM by addition of Retinal (RAL) to MG. Hallmark and KEGG datasets were employed. **e,** Expression changes compared to the control (without MG) samples are indicated for the different conditions (RAL or RA addition) at both 96h and 120h. Genes indicated here are DEGs identified in Fig.3g (96h MG). **f-h,** Expression levels of indicated genes across different conditions in SM for both 96h and 120h Axioloids.

**Extended Data Fig.9.**
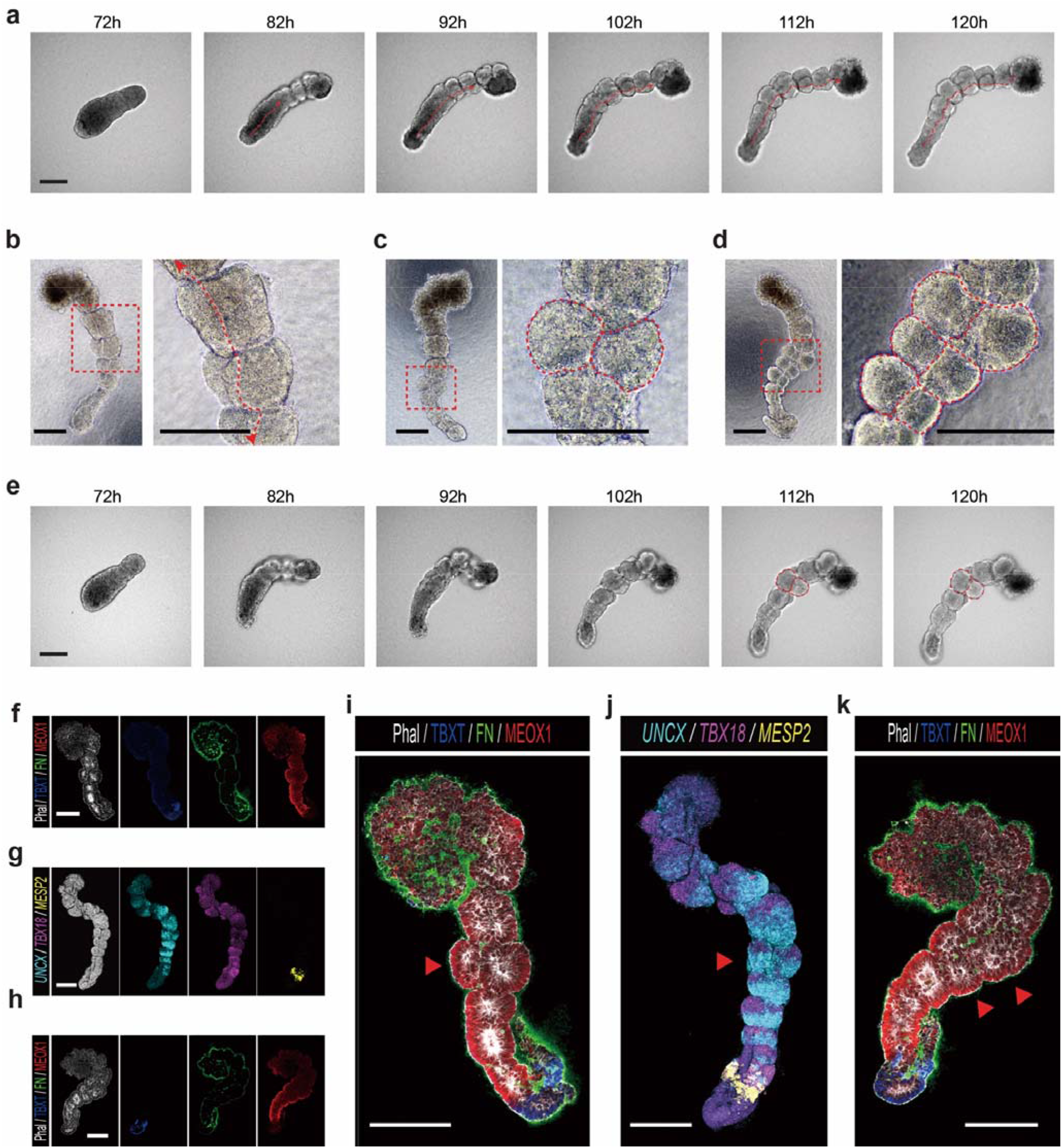
Midline and bilateral somite formation in human Axioloids. **a**, and **e**, Serial images of elongating human Axioloids (409B2) at 8-10h intervals, from 72h to 120h (extracted from **Supplementary Video 5**). **a**, Red dotted arrow shows formation and extension of superficial midline along the posterior to anterior axis of an Axioloid embedded in MG+RAL. **e**, Red arrowhead shows the initiation of somite division into 2 bilateral structures highlighted and circled with dotted lines, Axioloid embedded in MG+ROL. **b**-**d**, Left, brightfield images of Axioloids (409B2) at 120h, red dotted square encompasses the area enlarged in the right image. **b**, MG+RAL embedded Axioloid showing a noticeable midline highlighted by a dotted double arrow in the right image. **c**, and **d**, respectively MG+RAL and MG+RA embedded Axioloids showing in **c**, a single bilateral somite and in **d,** multiple bilateral somites highlighted by red dotted lines. **f**, **h**, **i**, **k**, Immunofluorescence staining of F-actine (Phalloidin) in gray, TBXT (BRA) in blue, FN in green and MEOX1 in red. **g**, and **j**, HCR staining of *MESP2* in yellow, *UNCX* in cyan and *TBX18* in magenta of human Axioloid (409B2) at 120h. Axioloids in **f**, and **g**, present a single bilateral somite and in **h,** and **k**, multiple bilateral somites. **i**, **j**, and **k** Correspond to an enlarged view of the merged channel images of **f**, **g**, and **h**, respectively. Scale bar is 200μm.

**Extended Data Fig.10.**
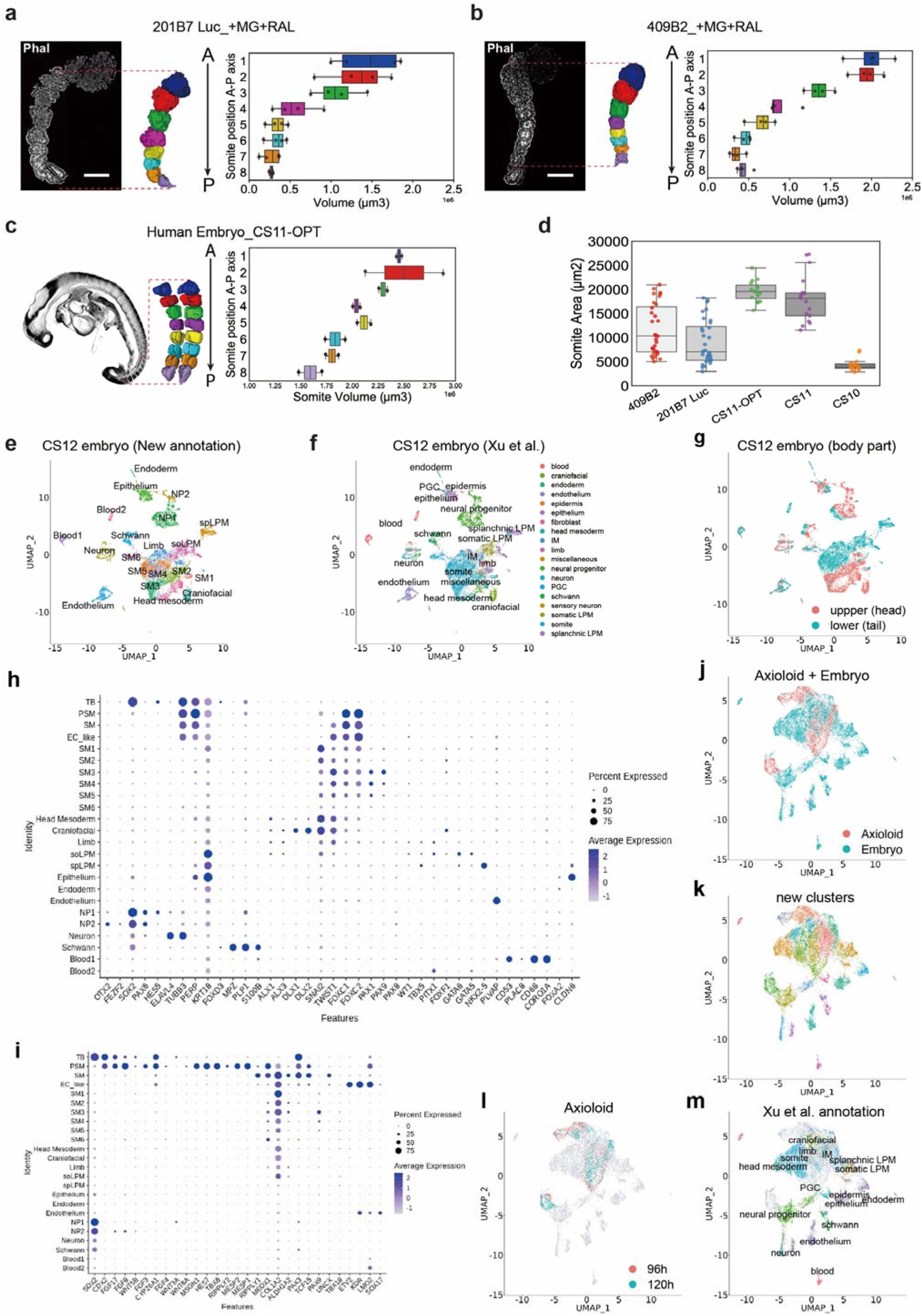
Morphometric and molecular (scRNA-seq) comparison of human Axioloids with human CS11 and CS12 embryos. **a,** and **b,** Phalloidin staining and z-stack image-based 3D model creation and somite volume measurement of **a**, 409B2 and **b**, 201B7 Luc human Axioloids embedded in MG+RAL at 120h (N=2, n=4). **c**, and **d,** OPT stack single image and image stack-based 3D model creation and somite volume measurement of the 8 posterior somites of a CS11 human embryo (Left and right side n=2, OPT data of human embryo obtained from the Human Developmental Biology Resource (HDBR)). **d**, Measurment of the somite area on histological section of CS11 (n=2) and CS10 embryo (n=1) and comparison with the somite area extrapolated from the volumes measured in **a-c**. **e**-**f**, UMAP projection of scRNA-seq datasets of CS12 human embryo. **e,** Colored based on our clustering **f,** on cell annotations by Xu et. al^35^ and **g**, on the sample origins. **h,** and **i,** Averaged expression levels of indicated genes in each cluster. **h,** Shown genes are marker genes of annotated clusters of Xu et. al and **i,** marker genes of Axioloids. **j**-**m,** UMAP projection of integrated scRNA-seq profiles of Axioloids and human embryos, colored by **j,** their origins (Axioloid or embryo), **k,** defined clusters **l,** Axioloid samples (96h or 120h) and **m**, cell types (annotated by Xu et al).

**Extended Data Fig.11.**
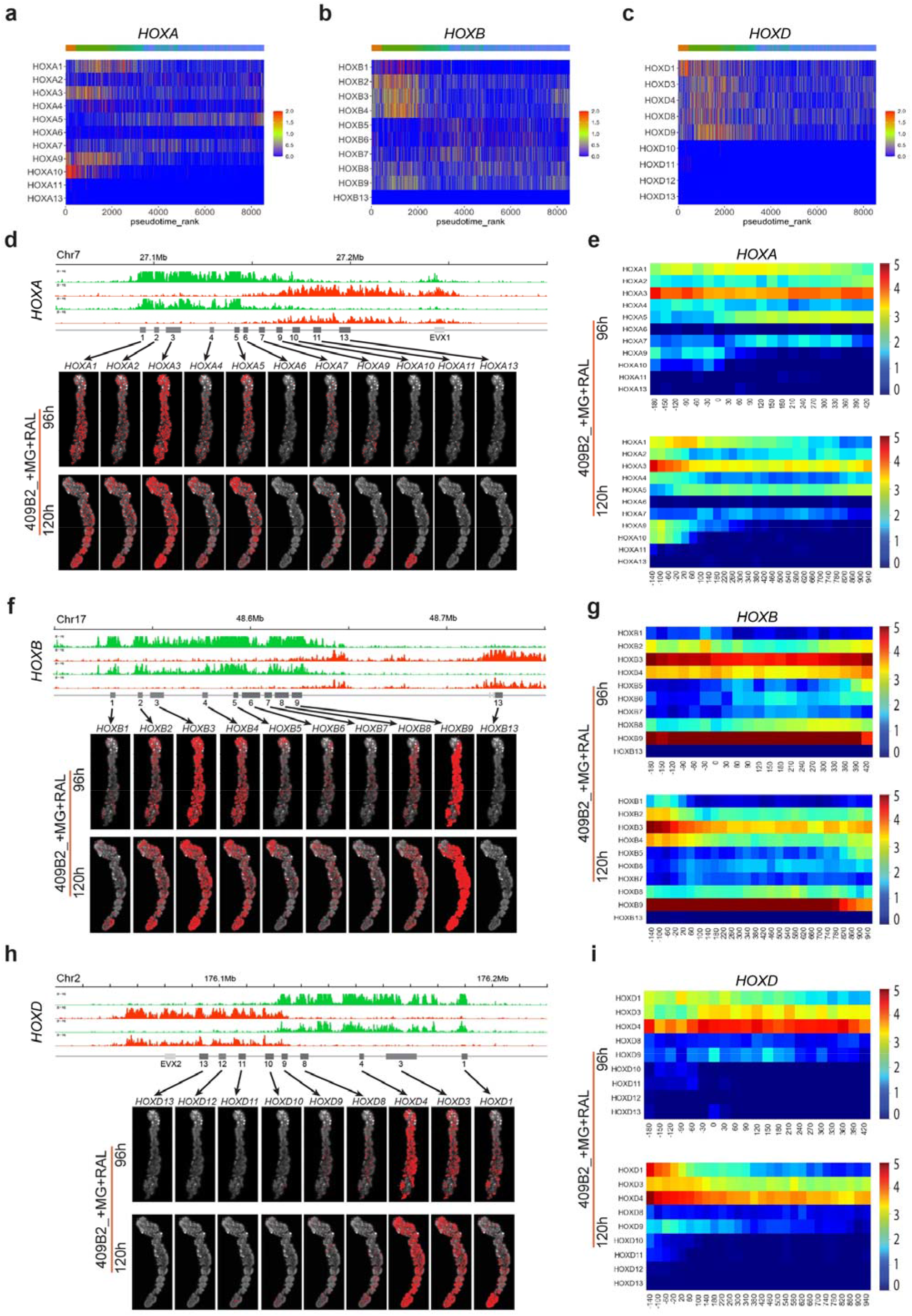
Assessment of the HOX Code in human Axioloids. **a-c,** Pseudotime representation of expression of HOXA, HOXB and HOXD cluster associated genes in human MG exposed Axioloids at 96h of culture (24h after embedding into MG); gene expression patterns arranged along pseudotime rank. **d, f,** and **h,** Top panels, analysis of the epigenetic landscape at the HOXA, HOXB and HOXD loci profiled by CUT&Tag using antibodies against H3K4me3 (green) and H3K27me3 (red). Bottom panels, visualization of the spatial distribution of the HOXA, HOXB and HOXD transcripts using HybISS analysis of all the members of the respective HOX clusters using sections of 96h and 120h Axioloids cultured in MG+RAL. **e, g,** and **i,** Heatmap plots of the HybISS data shown in d, f, and h, showing the average HOXA, HOXB and HOXD cluster gene expression along the posterior to anterior axis of 96h and 120h MG+RAL Axioloids normalized to the position of the *MESP2* signal peak (n=3).

**Extended Data Fig.12.**
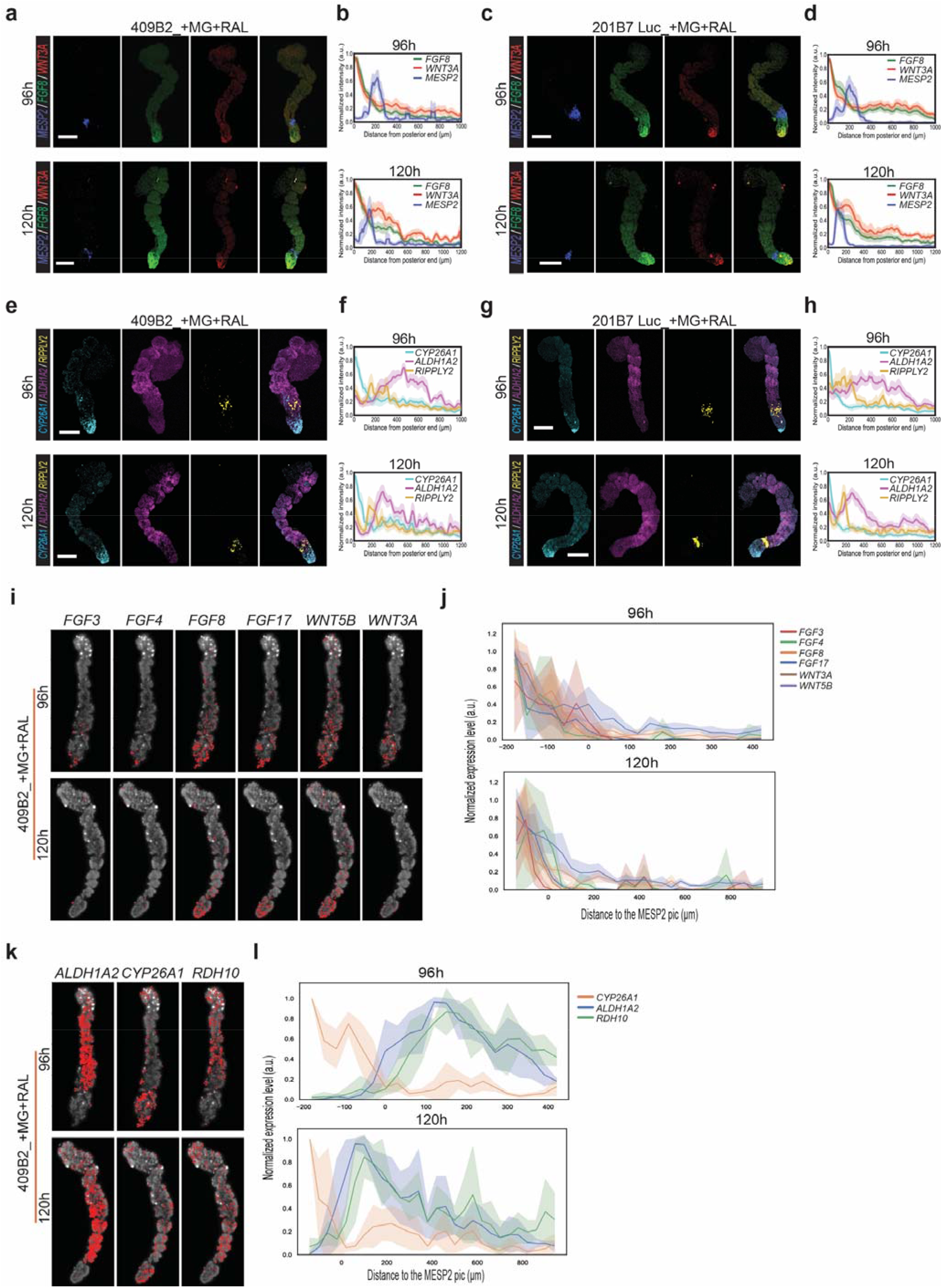
Expression gradients of FGF, WNT and RA signaling pathway members in human Axioloids visualized and quantified via HCR and HybISS. **a-h,** HCR whole mount in situ hybridization images and signal quantification of MG+RAL embedded Axioloids at 96h and 120h derived from **a, b, e, f,** 409B2 and **c, d, g, h,** 209B7 Luc iPSC lines. Shown images are representative of 3 independent experiments. **a,** and **c,** HCR staining of *FGF8* in green, *WNT3a* in red and *MESP2* in blue. **b,** and **d,** corresponding quantification along the posterior to anterior axis of the signal intensity normalized the position of the *WNT3a* signal peak (409B2 96h: N=4, n=7 and 120h: N=3, n=6, 201B7 Luc 96h: N=3, n=9 and 120h: N=3, n=8,). **e,** and **g,** HCR staining of *CYP26A1* in cyan, *ALDH1A2* in magenta and *RIPPLY2* in yellow. **f,** and **h,** corresponding quantification along the posterior to anterior axis of the signal intensity normalized to the position of the *CYP26A1* signal peak (409B2 96h: N=4, n=10 and 120h: N=3, n=7, 201B7 Luc 96h: N=3, n=8 and 120h: N=3, n=10,). Scale bar 200 μm. **i-l,** HybISS based visualization and quantification of the spatial distribution of FGF/WNT and RA signaling pathway transcripts in human Axioloids at 96h (top) and 120h (bottom) of culture in MG+RAL. **i,** and **j,** HybISS images and quantification of spatial expression of *FGF3, FGF4, FGF8, FGF17, WNT3a* and *WNT5b*.**k,** and **l,** HybISS images and quantification of spatial expression of *ALDH1A2, CYP26A1* and *RDH10*.

**Extended Data Fig.13.**
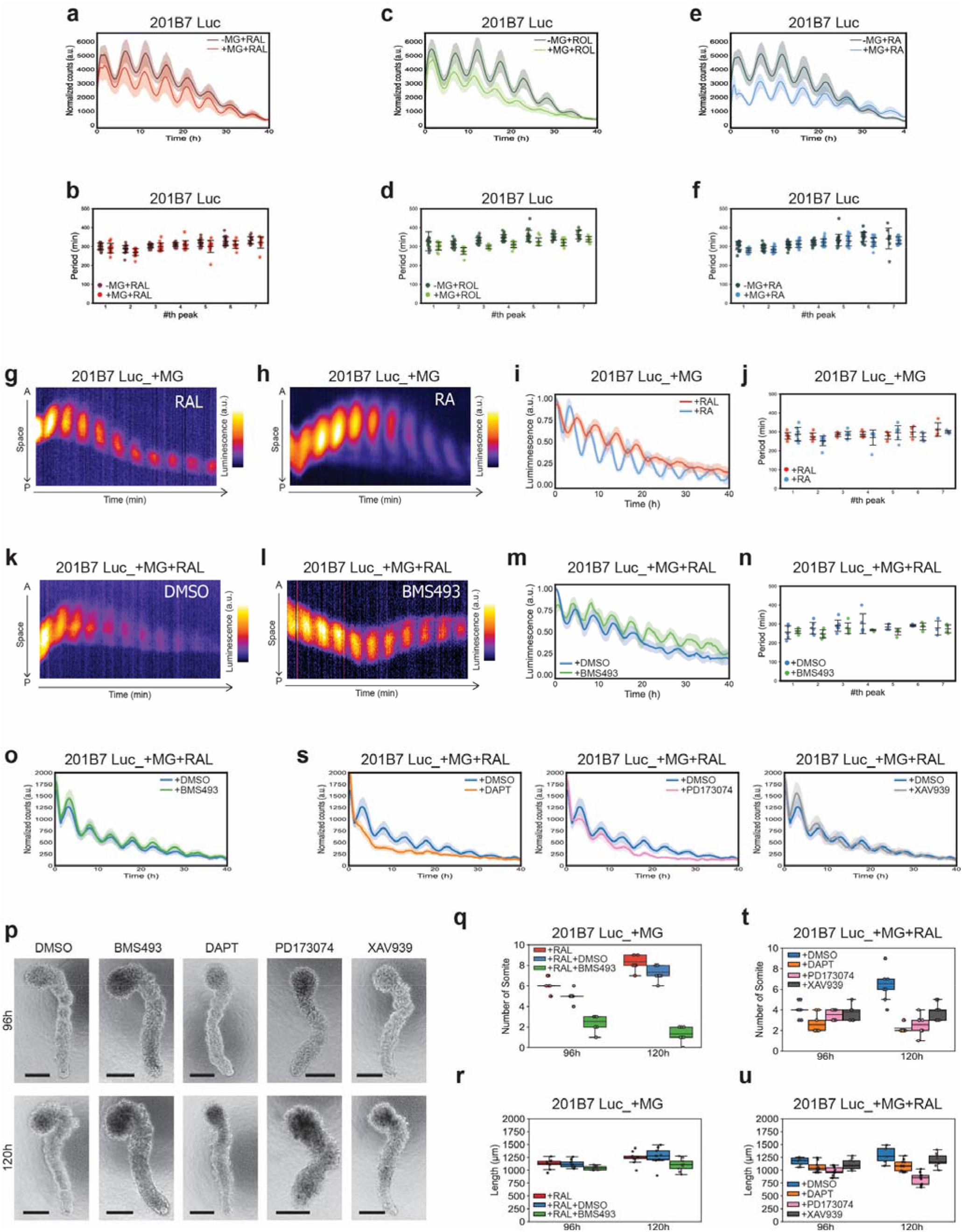
Modulating RA, FGF, WNT and Notch signaling in human Axioloids. **a-o** and **s,** Quantification of the total HES7:luciferase signal over time and corresponding period measurements as the time interval between the consecutive HES7:Luciferase signal peaks in Axioloids +/− MG in **a**, **b**, +RAL (Retinal) (N=5, n=17 and N=5, n=20), **c**, **d**, +ROL (Retinol) (N=3 n=12 and N=5, n=20), **e**, **f**, +RA (Retinoic Acid) (N=5, n=20 and N=5, n=20), **f**, (N=3, n=22 and N=5, n=20). Kymographs of the HES7:luciferase signal for Axioloids embedded in **g,** MG+RAL (N=4, n=6 data identical to the one showed in Fig.4 g, n, u), **h,** MG+RA ( N=4, n=8), and corresponding **i,** average signal and **j,** periodicity measurement, **k,** MG+RAL+DMSO (N=3, n=6), **l,** MG+RAL+BMS493 ( N=3, n=6) and corresponding **m,** average signal and **n,** periodicity measurement. **o,** Effect of the addition of BMS493 (N=3, n=9), **s,** of DAPT (N=3, n=9), PD173074 (N=3, n=9), and XAV939 (N=3, n=9), compared to DMSO (N=3, n=9) on the total HES7:luciferase signal over time in MG+RAL Axioloids. **p,** Representative brightfield images of Axioloids embedded in MG+RAL supplemented with DMSO or BMS493 or DAPT or PD173074 and XAV939, at 96h and 120h. **q,** Comparison of the total number of somites of Axioloids embedded in MG+RAL only or supplemented with DMSO or BMS493 or **t**, DAPT, PD173074, XAV939 (N=3, n=9 for all), and length of the Axioloids at 96h and 120h after addition of DMSO (N=3, with n=9) and BMS493 (N=3, n=9) in **r**, or DAPT, PD173074, XAV939 (N=3, n=9) in **u**. Scale bar is 200μm.

**Extended Data Fig.14.**
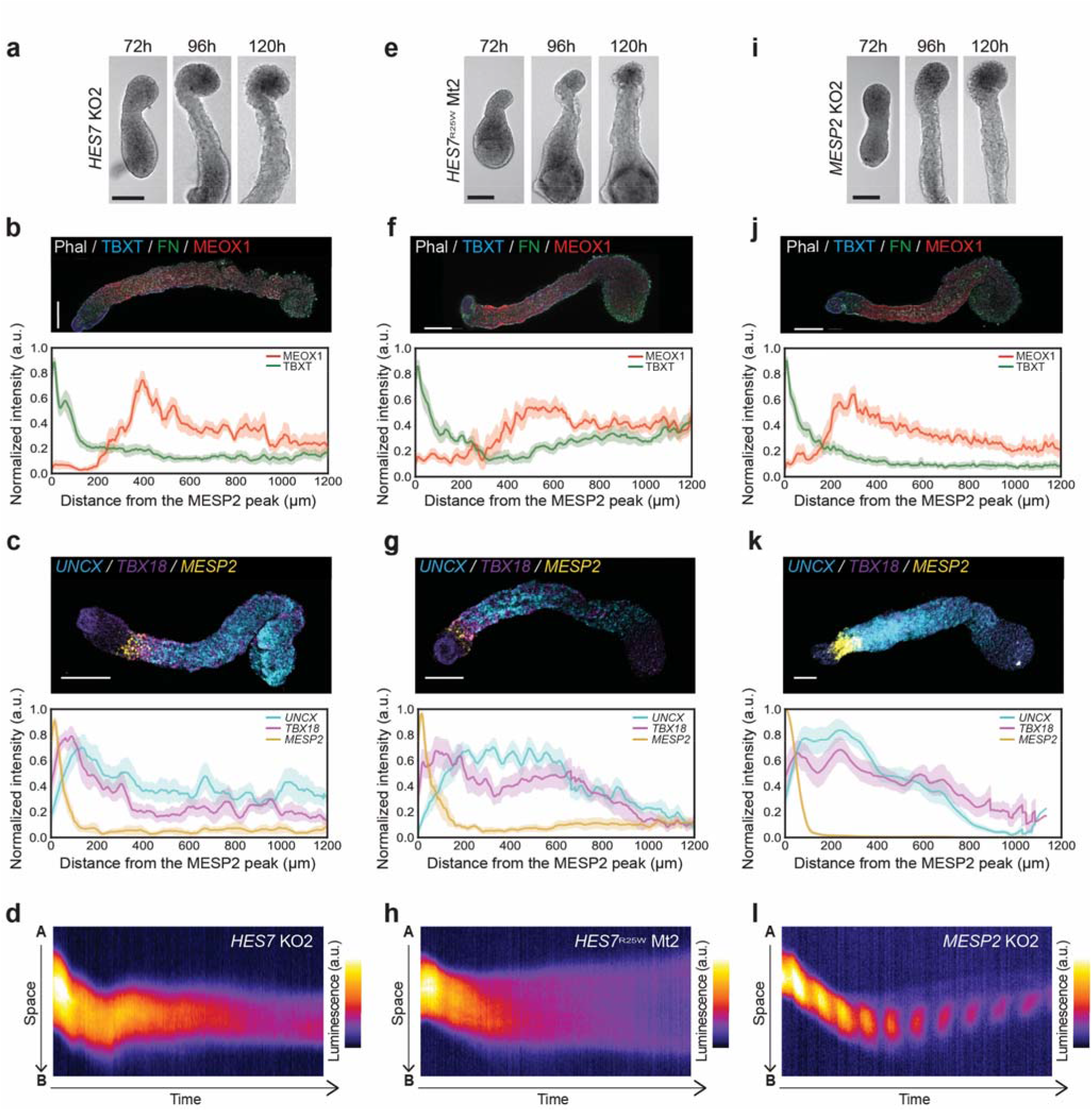
Morphological, molecular and functional characterization of patient-like iPSC-derived Axioloids with mutations in HES7 and MESP2. Panels **a-d,** show data for *HES7* KO2, panels **e-h,** show data for *HES7^R25W^* MT2 and panels **i-l,** show data for *MESP2* KO2. **a, e,** and **i,** Serial brightfield images of a forming Axioloid at 72h, 96h and 120h extracted from the **Supplementary Videos 9, 11,** and **13** respectively. **b, f,** and **j,** Immunofluorescence staining and signal quantification of MG+RAL embedded Axioloids at 120h. Top, merged channel images of Axioloids stained for F-actine (Phalloidin) in gray, FN in green, and MEOX1 in red, TBXT (BRA) in Blue. Bottom, corresponding quantification along the posterior to anterior axis of TBXT (green line) and MEOX1 (in red) signal intensity for *HES7* KO2 (N=3, n=9), *HES7*^R25W^ MT2 (N=3, n=9), *MESP2* KO2 (N=3, n=12). **c, g,** and **k,** HCR staining and signal quantification of MG+RAL embedded Axioloids at 120h, *MESP2* in yellow, *UNCX* in cyan and *TBX18* in magenta, and corresponding signal intensity measurements along the posterior to anterior axis normalized to the position of the *MESP2* signal peak for *HES7* KO2 (N=3, n=8), *HES7*^R25W^ MT2 (N=3, n=8), *MESP2* KO2 (N=3, n=9). **d, h,** and **l,** Kymograph of *HES7* oscillatory activity for **d,** *HES7* KO2, **h,** *HES7^R25W^* MT2, and **l,** *MESP2* KO2 Axioloids embedded in MG+RAL. Scale bar is 200 μm.

